# Gas5 regulates early life stress-induced anxiety and spatial memory

**DOI:** 10.1101/2023.09.27.559778

**Authors:** Dipanjana Banerjee, Sania Sultana, Sourav Banerjee

## Abstract

Early life stress (ES) induced by maternal separation (MS) remains a proven causality of anxiety and memory deficits at later stages of life. Emerging studies have shown that MS-induced gene expression in the hippocampus is operated at the level of transcription. However, the extent of involvement of non-coding RNAs in MS-induced behavioural deficits remains unexplored. Here, we have investigated the role of synapse-enriched long non-coding RNAs (lncRNAs) in anxiety and memory upon MS. We observed that MS led to an enhancement of expression of the lncRNA Gas5 (Growth Arrest Specific-5) in the hippocampus; accompanied by increased levels of anxiety and deficits in spatial memory. Gas5 knockdown in early life was able to reduce anxiety and partially rescue the spatial memory deficits of maternally separated adult mice. However, the reversal of MS-induced anxiety and memory deficits is not attributed to Gas5 activity during neuronal development as Gas5 RNAi did not influence spine development. Gene Ontology analysis revealed that Gas5 exerts its function by regulating RNA metabolism and translation. Our study highlights the importance of MS-regulated lncRNA in anxiety and spatial memory.

## INTRODUCTION

Maternal separation, a well-established model of early-life stress, has been extensively studied to investigate the long-term consequences of adverse early life events during this critical period of brain function and behaviour (Taylor et al., 2010; Korosi et al., 2012). Stress due to maternal separation (MS) in rodents leads to anxiety-like behaviours in adulthood as characterized by heightened fear, avoidance, and stress sensitivity (Banqueri et al., 2017; Lundberg et al., 2020; Balouek *et al*., 2023). The ventral hippocampus, a key brain area affected by MS (Dimatelis et al., 2014; Marais et al., 2009), has been shown to regulate emotions (Jimenez et al., 2018; H. Li et al., 2022; McNaughton, 1997). MS-induced anxiety can lead to changes in the total volume of the hippocampus (Kalisch et al. 2006) and alterations in the expression of glutamatergic receptors (Pickering et al. 2006), both NMDA (Barkus et al. 2010) and AMPA (Katsouli et al. 2014); GABAergic receptors (Sterley, Howells, and Russell 2013), and serotonergic as well as cholinergic receptors (File et al., 2000). Moreover, early-life stress can impact spatial memory that involves the dorsal hippocampus (Morris *et al*., 2006; Blum *et al*., 1999). Such behavioural deficits are linked with structural and functional modifications in the hippocampus (Izquierdo and Medina, 1997; Revest *et al*., 2009; Kremmyda *et al*., 2016). These observations indicate that the enduring effects of maternal separation on anxiety and memory involve alterations in synaptic plasticity (Monroy et al., 2010; Bannerman *et al*., 2014) and neurotransmitters, for example, noradrenaline and serotonin (Daniels et al. 2004).

More recently, transcriptomics analysis following MS identified a distinct set of miRNAs in specific brain areas including the hippocampus (Mckibben and Dwivedi 2021). Some of these miRNAs are localized at the synapse and have been shown to regulate synaptic plasticity and memory (Mckibben and Dwivedi 2021). A recent study from our laboratory has identified a set of synapse-enriched lncRNAs from the hippocampus. Among these transcripts, a specific lncRNA has been shown to regulate hippocampus-dependent memory (Samaddar et al., 2023). LncRNA functions as scaffolds, decoys, guides, or enhancers to modulate chromatin remodelling, transcription, mRNA stability, and RNA-protein interactions (Mattick et al., 2023; Mercer et al., 2009). Recent attention has focused on lncRNA activity at synapses that govern synaptic plasticity, neuronal development, and neurological disorders (Zhang, Hamblin, and Yin 2017) (Shi, Zhang, and Qin 2017) (A. Wang et al. 2017). The regulatory function of lncRNA has also been attributed to anxiety (Spadaro et al. 2015) and memory (Soutschek et al. and Schratt et al., 2023; Cui et al., 2022).

Prompted by these observations, we have investigated the impact of MS on the expression of lncRNAs and their involvement in MS-induced anxiety and memory. Our study identifies that MS enhances the expression of the lncRNA Gas5. The knockdown of Gas5 alone shows no significant changes in spine development in adult mice. However, when accompanied by early-life stress, Gas5 RNAi partially alleviates the enduring consequences of the stress as detected by the reduction in anxiety and the occlusion of memory deficits following MS.

## MATERIAL AND METHODS

### Animal ethics, acquisition, and maintenance

All animals were issued from the National Brain Research Centre (NBRC) animal house facility from the protocol numbers NBRC/IAEC/2015/171, NBRC/IAEC/2015/172 and NBRC/IAEC/2015/178 approved by IAEC (Institutional Animal Ethics Committee) of NBRC registered with the CPCSEA (Committee for the Purpose of Control and Supervision of Experiments on Animals) (Registration number: 464/GO/ReBi-S/Re-L/01/CPCSEA), Government of India.

Animals were housed in standard housing facilities in cages with food water *ad libitum*. A temperature of around 24± 0.5 °C and humidity of 40 ± 5% was maintained along with a 12-hour light/dark cycle for all animals.

### Maternal Separation

In our study, we have focused on the aspects of induction of stress only due to maternal separation and not early social isolation, where pups are separated from the mother as well as from each other (Alves et al., 2022). The maternal separation paradigm was used as a chronic early-life stress paradigm for C57BL/6 mice pups (Sood et al. 2018). A litter size of 5-7 pups was selected for the experiment. 1 litter of pups or 1 dam was considered ‘N’ and was arbitrarily allocated into homecaged (HC) and maternally separated (MS) groups. A total of 32 dams (120 pups (n)) were used for maternal separation throughout the study. Postnatal day 2 (P2) pups were separated from their mother for a period of 3 hours (0900 to 1200) till P14. The mother was first moved into a different fresh cage and kept in a separate room. The pups were then placed in a beaker with soft bedding and protective paper lining and the temperature was maintained at 37 ° C throughout separation. After 3 hours, first, the pups were kept back in the original cage followed by the mother. No separation was done from P15 till weaning on P21. Handling of the pups was done very gently. Cage changes were done every three days starting from P4. For the control group, designated as HC, all litters and mothers were kept undisturbed from P0 to P21 except for regular maintenance and cage changes every three days. Male pups were sacrificed at P21 (euthanization with CO_2_ gas), and hippocampi were isolated for further biochemical processes.

### Screening of Lnc RNAs

P21 male and female pups (N=14) were sacrificed in a CO_2_ euthanasia chamber at a 30-70% fill rate of CO_2_ gas and death was confirmed by tail/feet pinch and lack of respiration. Euthanasia was followed by a quick decapitation of the animals and the whole hippocampi was harvested in Trizol reagent (Invitrogen) for total RNA isolation. SuperScript III First Strand Synthesis System (Life Technologies, Invitrogen) kit was used for synthesizing cDNA following the manufacturer’s protocol. Further, qPCR of the cDNA prepared was done using SyberGreen (Applied Biosystems). See Table 1 for qPCR primer details.

**TABLE 1.**
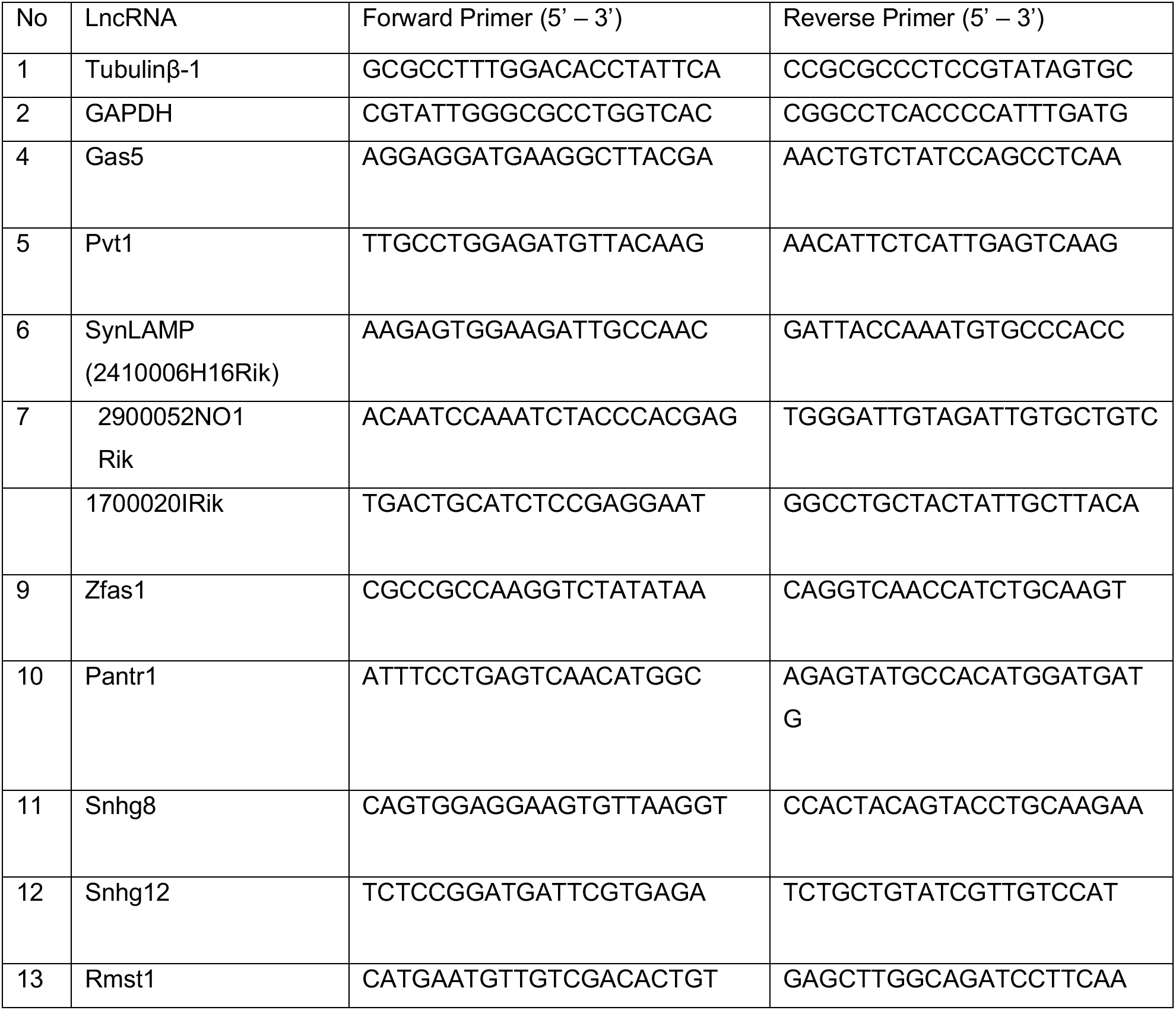
Primers for qPCR.

### Cloning and Virus preparation

EF1-α promoter from pLVTHM (Addgene # 12247) was used to replace the CMV promoter in pAAV.U6.shRLuc.CMV.ZsGreen.SV40 (Penn Core Vector) plasmid by excising out the CMV promoter using the EcoRI (NEB #R3101) and NHE-I (NEB #R3131) cut sites. Infusion cloning (Takara-Bio In-Fusion HD cloning Kit; 639650) was used to clone in EF1-alpha promoter generating a pAAV.U6.shRLuc.EF1-α.ZsGreen.SV40 plasmid and express ZsGreen under EF1-α promoter (Table 2). Universal Control (UC) and Gas5 shRNAs (shRNA # 1= G5C1 and shRNA # 2= G5C2) were cloned by replacing the existing Luciferase shRNA (shRLuc) under a U6 promoter using BamHI (NEB #R0136L) and EcoRI restriction sites (Table 3). The final vector was coined as pAAV.U6.shRNA(UC).EF1-α.ZsGreen.SV40, pAAV.U6.shRNA(G5C1).EF1-α.ZsGreen.SV40 and pAAV.U6. shRNA(G5C2).EF1-α.ZsGreen.SV40 respectively.

**TABLE 2.**
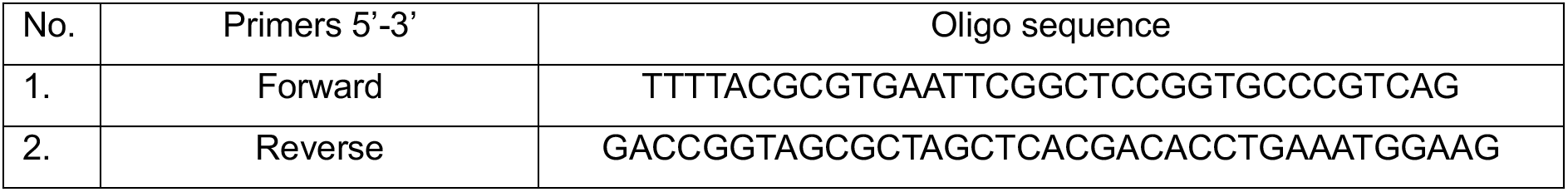
Infusion primers for replacing CMV with EF1-α in AAV backbone.

**TABLE 3.**
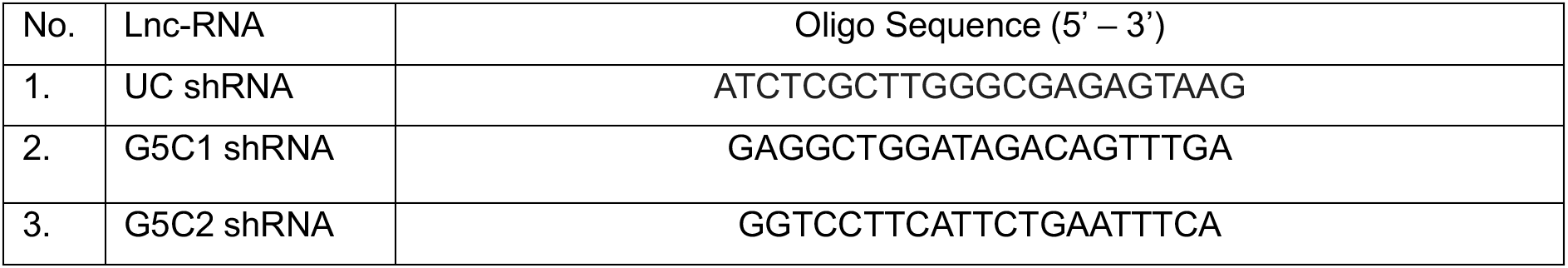
shRNA Antisense Sequence.

ZsGreen was replaced with EGFP in all vectors to achieve a diffused fluorescence for imaging. All vectors were digested with NcoI-HF (NEB #R3193) and NotI-HF (NEB #R3189) to excise out ZsGreen. EGFP from the pLVTHM vector was inserted by infusion cloning (Table 4).

All clones were verified by Sanger sequencing.

**TABLE 4.**
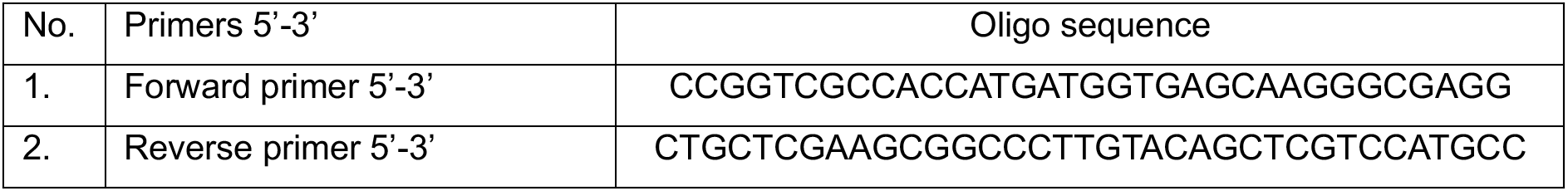
Infusion primers for replacing ZsGreen with EGFP in AAV backbone.

The expression vector was propagated using the *E.coli* Stbl3 strain and the AAV packaging plasmids pAdDeltaF6 and pAAV2.9 (Penn vector Core) were propagated using the DH5α strain. Endotoxin-free Maxiprep kit (Qiagen) was used to prepare purified plasmid for HEK293T cell transfection. Calcium phosphate transfection was done using 10µg of the above-mentioned transfer vector, 10µg pAAV2/9n and 20µg of pAdDeltaF6 into HEK293T cells of density 2X10^6^ cells. The culture was maintained at 37 °C, 5% CO_2_ in low-glucose DMEM (Gibco) with 10% foetal bovine serum with one-time media change after 8 hours. Post 96 hours of infection, NaCl (final concentration: 500 mM) was added to the plates for 3 hours to lyse the cells. The supernatant was collected and stored at 4 °C. The remaining cells on the plates were scraped and collected separately. The cells went through three freeze-thaw cycles where they were frozen at −80 °C and thawed at 37 °C for complete dissociation of virus particles from the cell membrane. The filtrate was added to the supernatant collected before; together they were filtered through a 100µ and a 0.45µ filter successively. The final filtrate, containing the virus, was then centrifuged using Amicron-Ultra-15 (Merk Millipore #UFC910024) filters to concentrate the virus and determine the titre by fluorescence-based cell counter. The multiplicity of infection (MOI) was calculated by infecting 80K HEK293T cells with serially diluted AAV solutions. An MOI of 1-2 was used for further experiments.

### Primary Neuronal Culture

Hippocampal neurons were cultured following the previous protocol (Kaech & Banker, 2006). CD1 dams were euthanized in a CO_2_ gas chamber (fill rate 30-70%) followed by cervical dislocation and death was confirmed by tail/feet pinch. The uterus containing the embryonic day 16 (E16) pups was dissected and taken out on ice-cold dissection media (Gibco) to render the pups unresponsive to painful stimuli. An incision was made in the uterus to extract the pups for decapitation and the whole brain was collected from the sacrificed pups. Hippocampi from all pups from a single CD1 (N=4) were pooled for further experiments. Briefly, hippocampi from embryonic pups were dissected and collected in dissection media (Gibco). It was then digested with the addition of 2.5% trypsin to the dissection media at 37 °C for 10 mins. Trypsin-treated tissue was washed with dissection media and collected in Glial Minimum Essential Media (GMEM; Gibco). Further, trituration was done with a fire-polished Pasteur pipette to obtain a single-cell suspension which was plated on poly-L-lysine coated 60mm dishes (1 mg/mL, Sigma) at a concentration of 200-250 cells/mm^2^. The culture was maintained at 37 °C, 5% CO_2_ in Neurobasal Media (Gibco) with B27 supplement (Gibco) throughout the experiment. Cells were infected with AAV at DIV 7 and harvested at DIV 21 for further experiments.

### Stereotaxic injection

Surgery was done in P0 pups (N=56, n=168) 6-8 hours post-birth only after the appearance of the milk spot (Kim et al. 2014). Half of the litter was taken at a time and kept on a heating pad at 37 ° C. For anaesthesia, the pup was kept on a cold metal plate kept on the ice for 2-3 mins till there was no movement and the body colour turned whitish pink from reddish pink. It was then lightly taped to the platform with its head erect. The platform was kept at 7-8 °C to maintain the anaesthesia. In case the pup’s body colour started to show a bluish tinge, the temperature was raised to 12-15 °C. Stereotaxic arms were moved to (X, Y) = (0.8, 1.5) mm coordinates and the needle was lowered till at Z= −1.8 mm (Christensen et al. 2010). Each hippocampus was injected with 0.5 µl of AAV with MOI=1. A volume of 0.1 µl was injected per minute with a gap of 10 mins. On completion of 0.5 µl virus injection, an additional gap of 10 mins was given before the needle was pulled out very slowly to prevent any back suction. We proceeded to inject into the second hippocampus after 15 minutes during which the first wound site was allowed to close. The pup was then kept on a 37 °C heating pad and warmed till movement and colour were regained. They were returned to the mother together to minimise disturbances. The cage was left completely undisturbed for the next day. (n=163; surgeries performed for western blot, spine analysis, light dark assay, spatial object recognition assay. ‘n’ determines the number of individual animals; males).

### Light/dark box assay

The apparatus consists of a smaller dark box with a light intensity of 2 Lux (one-third of the total area) and a bigger light box with a light intensity of 390 Lux (two-thirds of the total area) separated by a door (Bourin and Hascoët 2003) (Takao and Miyakawa 2006). P42 mice (N=26, n=79) were habituated for 2 minutes to handling and to a transport vessel in dim light (2-5 Lux) for 1 week in the experimental room. Animals were housed in standard cages and a 12-hour light/dark cycle was maintained as usual. On test day, the mouse was kept in the dark box with a closed door. The door was opened 10 seconds later and the mouse was left free to explore the light box for 10 minutes. The behaviour assays and habituations were performed between 0900 hours to 1800 hours. Arenas were cleaned using 70% alcohol between tests to prevent any odour-related bias (Takao and Miyakawa 2006). The video was recorded using a Logitech HD webcam C270 overhead-mounted camera and analysed using online free software (Toxtrac) (Rodriguez et al. 2018), which was also blind tested manually. The total distance covered, time spent in the light box, and the number of transitions made between the two boxes were considered. The litter effect for animals used in light/dark box assay was also analysed (Figure S 3A-C).

### Spatial object recognition assay

All animals were housed in single cages a week before the experiment. P50 male (N=27, n=84) C57Bl/6 mice were habituated for 2 minutes to handling and a transport vessel for 1 week followed by spatial object recognition assay which comprised a total of 5 sessions, each 10 minutes long. The process is similar to the classical object recognition assay with a few variations (Wimmer et al. 2012). A light intensity of around 15 Lux was maintained at the centre of the arena (Leger et al. 2013). Session 1 involved introducing the animal to the empty arena (12 cm X 15 cm with spatial cues) for short habituation to the open field. In sessions 2 to 4, the animal was trained to explore two objects of different shapes kept in their fixed coordinates, with a 5 cm gap between wall and object, through all the sessions. The animal was returned to its original cage for 5 minutes to relieve stress and the arena and objects were wiped with 70% alcohol between each session. The test session (5^th^ session) was carried out after 24 hours of the training sessions when only one object was moved to a new coordinate with leu to the spatial cues and the animal was allowed to inspect for 10 minutes. All recordings were done using a Logitech HD webcam C270 overhead-mounted camera and a MATLAB-based autotyping software was used to calculate the number of bouts on each object (T. P. Patel et al. 2014), which was also blind tested manually. A bout was considered when the animal moved its nose within a space 2 cm radially from the object. Any attempt to climb or jump on the object was not considered a bout. The Discrimination Index (DI), which is the ratio of the difference in time spent with either object to the total time spent with both objects during the test session, was calculated to verify the ability of the animal to successfully recognize the displaced object (Denninger, Smith, and Kirby 2018).

DI= [time spent with object displaced-time spent with not displaced]/ [time spent with displaced+ time spent with not displaced]

The litter effect for animals used in spatial object recognition assay was also analysed (Figure S 3D).

### Cryo-sectioning

Following behaviour assessments, animals underwent sacrifice at P60 (N=53, n=163). Mice were sedated with a heavy dose of Ketamine (60mg/kg) and Xylazine (10mg/kg) cocktail injected intraperitoneally. They were perfused intracardially with a solution of 1XPBS (phosphate buffer saline) followed by 4% paraformaldehyde, for maintaining tissue integrity. To ensure thorough fixation, brains were extracted and immersed in 4% paraformaldehyde for an additional 5 days post-perfusion. Subsequently, the fixed brains were incubated in 40% sucrose solution for a week till they settled to the bottom, enabling optimal cryo-sectioning. Brains were flash-frozen in dry ice and coronal sections of 50µ thickness were obtained and mounted on slides pre-coated with gelatin. The tissue sections were washed 3 times with autoclaved Mili-Q water and mounted with Vectasheild DAPI to stain the nucleus and visualized under the microscope.

### Spine analysis

To measure neuronal development, recombinant AAV expressing two distinct shRNAs were injected into the hippocampus of male mice (P0) (N=3, n=5) and these neurons were imaged at P38-P40 (16-bit) using a Nikon A1 HD25 point scanning confocal microscope with Nikon Plan Apochromat NA=1.4 oil immersion objective at 1024X1024 pixel resolution at 0.75X. 2X optical zoom was used to get high-magnification images of neurons with 10 −15 optical sections with 0.25µm step size. The GFP signal was captured by a 488 nm solid-state laser. Images were obtained under uniform parameters of laser power; pinhole diameter and detector gain across all conditions (Figure 6A-B).

Neurolucida 360 (MBF Bioscience) was used for spine morphology analysis. A 3-D skeletal image was generated using the Directional Kernel tracing algorithm and spines were detected and classified. Spine density was obtained by using Neurolucida Explorer (Figure 6C-E).

Coronal sections of the mouse brain were visualized using a Leica LAS V4.2 fluorescence microscope and Zeiss Apotome3 (magnified image) (Figure S 1G-H). All imaging parameters were kept uniform across experiments. Images were stitched post-acquisition using Fiji image analysis software (MosaicJ plugin).

### Western blot

The hippocampal tissue was obtained from C57BL/6 male P21 pups (N=8, n=25), previously subjected to injections of UC, G5C1, or G5C2 shRNAs at P0. The pups were euthanized in a CO_2_ chamber at a 30-70% fill rate of CO_2_ gas and death was confirmed by tail/feet pinch. The tissue was lysed in RIPA buffer (composed of 150mM NaCl, 0.1% NP-40, 0.5% sodium deoxycholate, 0.1% SDS, 50 mM Tris, pH 7.4) and subsequently centrifuged at 4°C for 10 minutes at 4000 g to remove debris. Protein quantification was performed using the Pierce BCA Protein Assay Kit (Thermofisher) with the resulting clear supernatant.

For experimental purposes, specific sample lysates were mixed with Laemmli Buffer and heated to 95°C. An amount of 5 µg of each sample was utilized for housekeeping gene tubulin-β1 (Tuj-1) blot, while 30 µg of each sample was employed for pan-Ago blot. The samples were resolved by electrophoresis on a 10% polyacrylamide gel and then transferred onto a nitrocellulose membrane. Subsequently, the membrane was blocked with 5% BSA and incubated with primary antibodies against Tuj-1 (dilution 1:10,000, Sigma, Catalog # T8578) and pan-Ago (dilution 1:1200, MerkMillipore, Catalog # MABE56) overnight at 4°C. Following the primary antibody incubation, the membrane was washed 6 times, 10 minutes/wash, with 1X TBST (Tris-buffered saline with 1% Tween 20) at RT. The blot was then exposed to secondary antibodies (dilution 1:10,000 for Tuj-1 and 1:5000 for pan-Ago, Goat Anti-Mouse, Thermofisher Scientific, Catalog# A11030) for 3 hours. Further washing was done with 1X TBST (6 washes, 10 minutes/wash) to obtain cleaner background. Luminescence on the blot was detected using the ECL chemiluminescence kit (Millipore). Band intensities and the expression of pan-Ago were quantified using ImageJ software, with normalization against tubulin-β1.

### Statistics

Statistical analyses were performed for all experiments. Data shown are represented as mean±SEM. The number of pregnant dams/ litters used is defined by ‘N’ and the total number of pups (obtained from the dams) used is designated as ‘n’. The number of neurons considered for spine analysis is represented as ‘*n’*’. qRT-PCR data for lncRNA screening, Argonaute (Ago) immunoblots and spine density were analyzed using unpaired Student’s t-test with Welch’s correction for unequal variance. qRT-PCR data for Gas5 knockdown efficiency were analyzed using paired Student’s t-test. Two-way ANOVA with Fisher’s LSD test was used for analyzing light/dark and SOR data analysis. GPower 3.1 was used for post-hoc power analysis for all experiments. The outlier test was done using the ROUT method (0.1% aggressive). Immunoblot band intensity analysis and image stitching were done using ImageJ software. Gene Ontology plots were computed using RStudio (Figure 5 A-C) and SRplot ( Figure 7 B-E) (Tang et al. 2023). Data with a p-value less than 0.05% were considered significant. All data were analysed using GraphPad Prism 9. No randomisation technique was used animals were grouped arbitrarily. No prior sample size was calculated. Post-hoc power analysis was done for all experiments. Detailed statistical analysis and criteria for the exclusion of animals is provided in Supplementary data and Figure S5.

## RESULTS

### Maternal separation alters the differential expression of synapse-enriched lncRNAs

To investigate the impact of early life stress, this study used the maternal separation (MS) paradigm (Figure 1A) and the differential expression of a subset of synapse-enriched lncRNAs was analyzed by qPCR in males and females separately. In males, we observed a significant upregulation in the expression of specific lncRNAs, such as Pantr1 (146.3 ± 56.78% increase, p=0.041) and Gas5 (79.6 ± 19.14% increase, p=0.006) and without any change in expression of 1700020I14Rik (8.30 ± 13.14% increase, p=0.5522) upon MS. Although, we observed an increase in expression of 2410006H16Rik (105.5 ± 75.28% increase, p=0.210), 2900052NO1Rik (100.3 ± 70.88% increase, p=0.206), Pvt1 (82.9 ± 46.95% increase, p=0.127), RMST_1 (49.9 ± 51.05% increase, p=0.366), Snhg6 (95.4 ± 81.12% increase, p=0.284), Snhg8 (80.5 ± 73.63% increase, p=0.316), Snhg12 (86.3 ± 48.60% increase, p=0.126) and Zfas1 (37.4 ± 23.57% increase, p=0.163); these enhancement in expression are not statistically significant (Figure 1B). Gas5 was selected as the target lncRNA and two shRNAs (G5C1 and G5C2) against Gas5 were cloned into an AAV vector (Figure S1A). Knockdown efficacy was assessed in primary hippocampal neuronal culture and successful knockdown was evaluated for both the constructs (82.82 ± 9.33 % decrease, p= 0.003 for G5C1 and 84.93 ± 13.68 % decrease, p= 0.25 for G5C2 w.r.t GAPDH; 81.3 ± 8.46 % decrease, p=0.0024 for G5C1 and 83.71 ± 6.9 % decrease, p=0.0067 for G5C2 w.r.t Tubulin-β1) (Figure S1C-D).

**Figure 1.**
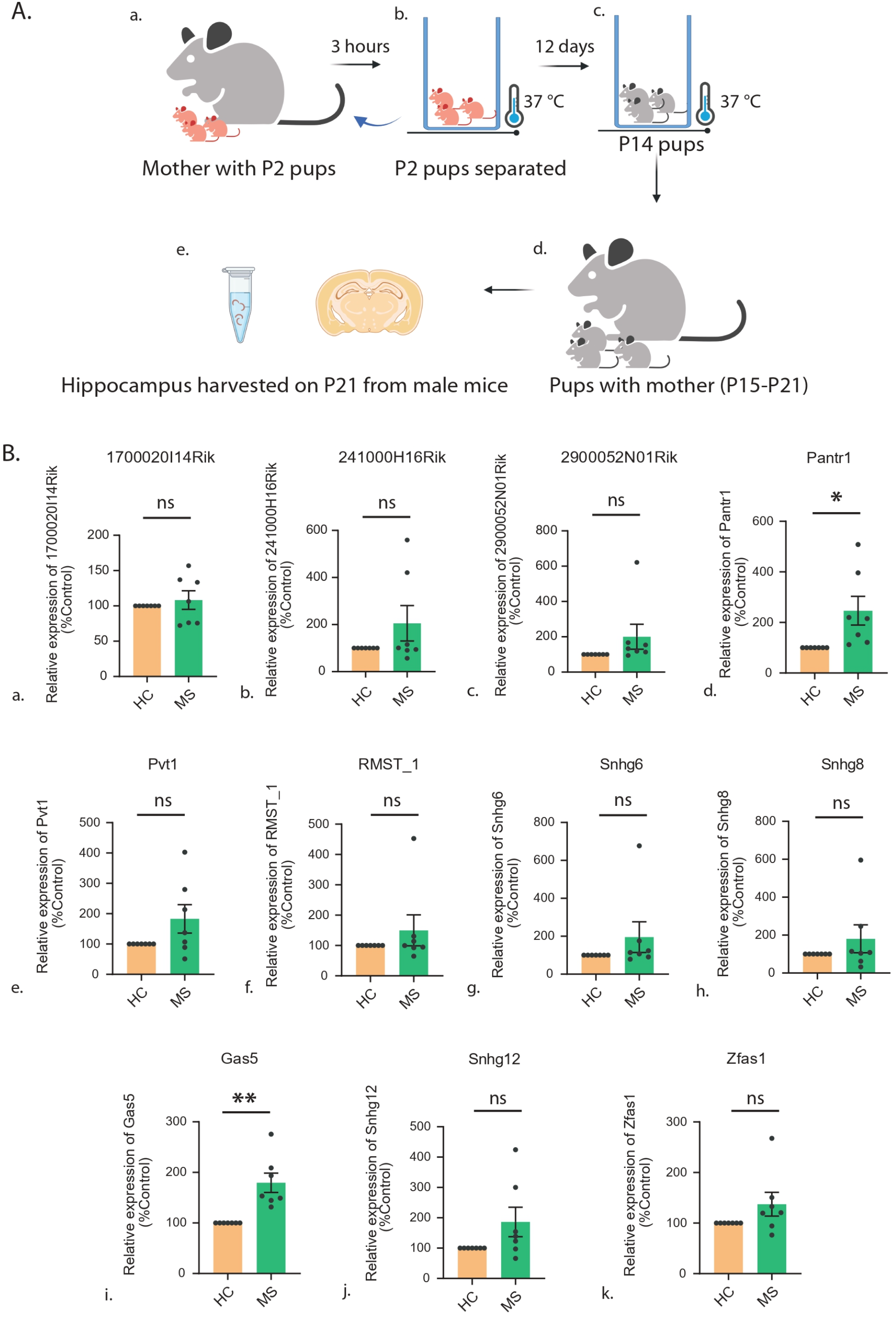
Maternal separation affects the expression of synapse-enriched lncRNAs. (**A)**. Maternal separation paradigm followed: P2 pups separated from their mother for 3 hours during P2 – P14 and following MS pups were kept till P21. (**B).** Differential expression of synapse-enriched lnc-RNAs in male hippocampus (P21) (a) 1700020I14Rik, (b) 2410006H16Rik, (c) 2900052NO1Rik, (d) Pantr1, (e) Pvt1, (f) RMST_1, (g) Snhg6, (h) Snhg8, (i) Gas5, (j) Snhg12, (k) Zfas1 as obtained through qPCR, *p < 0.05, **p < 0.01, ns = not significant. N=7, Data shown as mean ± SEM. Unpaired Student’s t-test with Welch’s correction. The underlying data is also shared in https://figshare.com/articles/dataset/_b_Gas5_regulates_early_life_stress-induced_anxiety_and_spatial_memory_b_/26047969.

In females, no significant alteration of Gas5 expression was seen after MS (24.77 ± 21.91, p= 0.3095). A significant downregulation of Zfas1 (47.50 ± 5.535, p= 0.0004) was observed. 2900052NO1Rik (164.9 ± 54.88, p= 0.0300), Snhg6 (101.9 ± 39.48, p= 0.0494) and Snhg8 (19.09 ± 6.565, p= 0.0335) were significantly upregulated upon MS. Other lncRNAs like 1700020I14Rik (5.670 ± 10.08, p= 0.5982), 2410006H16Rik (−28.74 ± 16.25, p= 0.1373), Pantr1 (−13.13 ± 18.85, p= 0.5172), Pvt1 (−20.40 ± 18.66, p= 0.3241), RMST_1 (219.2 ± 210.8, p= 0.3460) and Snhg12 (−5.823 ± 37.76, p= 0.8835) were not differentially regulated in the hippocampus after early life stress. The data showed the sexual dimorphic expression of lncRNAs between males and females.

### Gas5 knockdown partially alleviates MS-induced anxiety

We have analysed open-space anxiety of mice that are injected with AAV expressing shRNAs against Gas5 or non-targeting control. Mice were bilaterally injected with AAV expressing either G5C1 or G5C2 in the hippocampus at P0. These mouse pups were maternally separated or homecaged (P2 – P14); following which they were kept with their respective mother till P21 (day of weaning). Young-adult male mice (P48) were subjected to the light/dark box assay (Figure 3A) (X. Wang et al. 2022). Maternally separated mice exhibited avoidance of brightly-lit open areas as detected by subsequent reduction in time spent (102.2 ± 24.35 second (s) decrease, p<0.0001); distance covered (18.26 ± 2.814 meters (m) decrease, p<0.0001), and fewer transitions (13.55 ± 3.55 decrease, p= 0.0003) into the brightly lit arena as compared to the homecaged control. However, the maternally separated Gas5 RNAi mice showed a partial rescue in anxiety level as measured by time spent (65.47 ± 27.03 s increase, p= 0.0179 for G5C1 and 66.62 ± 25.21 s increase, p<0.0101 for G5C2), distance covered (17.45 ± 3.123 m increase, p <0.0001 for G5C1 and 13.38 ± 2.912 m increase, p<0.0001 for G5C2) and number of transitions (11.42 ± 3.943 increase, p= 0.005 for G5C1 and 8.615 ± 3.677 increase, p=0.0219 for G5C2) into the brightly lit arena with respect to maternally separated mouse expressing non-targeting control shRNA (Figure 3C-E). We have also observed that homecaged mice expressing Gas5 shRNA did not show any change in time spent, distance covered and the number of transitions into the brightly lit arena as compared to homecaged mice expressing non-targeting shRNA (Figure 3C-E). These observations indicate that Gas5 RNAi does not alter any physiological phenotype of the animal. Litter effect was also statistically analysed and no significant difference in behaviour among the litters was found (Figure S 3 A-C).

### Gas5 RNAi rescues MS-induced spatial memory deficit

We have analysed the impact of early-life stress on memory using spatial object recognition (SOR) test and evaluated the role of Gas5 following its knockdown (Figure S1A). Mice were injected (P0) with AAV expressing shRNA against Gas5 or non-targeting control. These mice were maternally separated or homecaged (P2 – P14) and subjected to SOR (P56). Mice were allowed to explore two distinct objects placed in two specific locations within an arena during the training session (Figure 4A, S2). On the test day, 24 hours after training, one of the objects was displaced within the same arena and the exploration time of non-displaced and displaced objects was analysed (Figure 4A-3B). We observed that the maternally separated mice expressing non-targeting shRNA displayed reduced recognition of the displaced object during the test session as determined by the discrimination index (DI) (0.4152 ± 0.1204 decrease, p=0.0009) when compared to the homecaged control (Figure 4D). Next, we examined the effects of maternal separation on memory following the knockdown of Gas5. We observed that the maternally separated mice having Gas5 RNAi could partially enhance DI (0.5568 ± 0.116 increase, p<0.0001 for G5C1 and 0.5186 ± 0.1181 increase, p<0.0001 for G5C2) as compared to the maternally separated mice injected with non-targeting control shRNA. Also, we have detected comparable DI of homecaged mice injected with AAV expressing either Gas5 shRNAs or non-targeting control shRNA (Figure 4D). Furthermore, the exploration of objects that were not displaced within a separate arena remains unaffected following Gas5 RNAi and maternal separation as we did not observe any change in DI across experimental conditions (Figure 4C). These observations indicate that the reduction of Gas5 expression could partially rescue deficits in spatial memory following maternal separation. Litter effect was also statistically analysed and no significant difference in behaviour among the litters was found (Figure S 3D).

### Gas5 is potentially involved in regulating RNA metabolism

To gain a plausible mechanistic insight into the Gas5-mediated regulation of anxiety and memory upon maternal separation, we took an *in silico* approach to identify the expression of transcripts influenced by Gas5. First, we have selected Gas5 interacting coding transcripts from the available CLIP-seq data (POSTAR3 database – http://postar.ncrnalab.org./) (Hu et al. 2017). We then performed a Gene Ontology (GO) analysis of these interacting coding transcripts and focused on significant enrichment (FDR < 0.05) of transcripts in specific GO terms (Figure 5A-C). The cellular processes included enrichment of GO-terms for ribonucleoprotein complex (FRD= 1.05E-31), cytoplasmic stress granule (FDR= 1.41E-17), polysome (FDR= 3.50E-05), somatodendritic compartment (FDR= 0.010067571) and dendrite (FDR= 0.004478929) (Figure 5B) etc. The molecular functions are enriched with GO-terms representing ribonucleoprotein complex binding (FDR= 1.22E-05), mi-RNA binding (FDR= 6.05E-17) and translation regulator activity (FDR= 7.53E-05) (Figure 5C). The biological process displays significant enrichment for GO-terms, such as RNA processing (FDR= 1.58E-43), RNA splicing (FDR= 8.08E-45), regulation of translation (FDR= 2.52E-18) etc. alongside some not significant but enriched terms like dendritic transport of ribonucleoprotein complex (FDR= 0.094670904) and miRNA transport (FDR= 0.063158655) (Figure 5A) (For details see Table S1 and Supplementary data). Overall, the GO analysis revealed that Gas5 is primarily involved in RNA metabolism including stability and regulation of translation. Prompted by this observation, we focused on the transcript encoding Argonaute (Ago) family of proteins that are involved in translation and RNA stability. As a proof-of-concept, we have analyzed the expression of Ago following Gas5 knockdown. Mice were injected with AAV expressing shRNA against Gas5 or UC (P0), and Ago expression in the hippocampus was analyzed at P21. We observed a significant increase in Ago expression in G5C1 (0.88 ± 0.25 increase, p=0.0084) and G5C2 (0.78 ± 0.29 increase, p=0.0335) injected animals as compared to Control RNAi individuals (Figure 5 D-E).

### Gas5 does not regulate spine development

To analyse the importance of Gas5 in spine development, we effectively knocked down Gas5 using two shRNAs at P0 and studied the changes in spine density at P30. We studied the spine density in neurons of the CA3 region of the hippocampus. No significant difference was observed in total spine density between the Control RNAi and either of the Gas5 RNAi neurons (0.01281±0.05731, p= 0.825 for G5C1 and 0.02916±0.06329, p=0.649 for G5C2) (Figure 6C). Additionally, we observed no significant difference in the density of stubby and mushroom spines between the Control RNAi group and the Gas5 RNAi individuals (Stubby: 0.0029± 0.034, p= 0.931 for G5C1 and 0.0533±0.034, p= 0.138 for G5C2; Mushroom: 0.0232±0.0171, p= 0.1869 for G5C1 and 0.0265±0.0177, p= 0.1504 for G5C2) (Figure 6D-E).

## DISCUSSION

LncRNAs emerged as key regulatory RNAs involved in diverse cognitive functions (Mercer, Dinger, and Mattick 2009) (Samaddar and Banerjee 2021) (Liau et al. 2021). A recent study from our laboratory identified a set of synapse-enriched lncRNAs from the hippocampus (Samaddar et al., 2023). This study highlighted the importance of one such synapse-enriched lncRNA, Gas5, which in early life stress, evoked behavioural deficits. We observed that early life stress of maternal separation induces the expression of Gas5 in the hippocampus of male mice (Figure 1B). The reduction of Gas5 expression by RNAi alleviates MS-induced anxiety and rescues memory deficits (Figure 3–4). Apart from Gas5, maternal separation also induces the expression of another synapse-enriched lncRNA, Pantr1 (Figure 1B). The hippocampi of female pups (P21) were collected similarly to the male pups. The screen revealed a very different regulation of lncRNAs in female pups upon MS revealing the sexually dimorphic outcomes due to stress. Further behavioural experiments were done in males only as female behaviour is evidenced to be influenced by the oestrous cycle (Figure 2).

**Figure 2.**
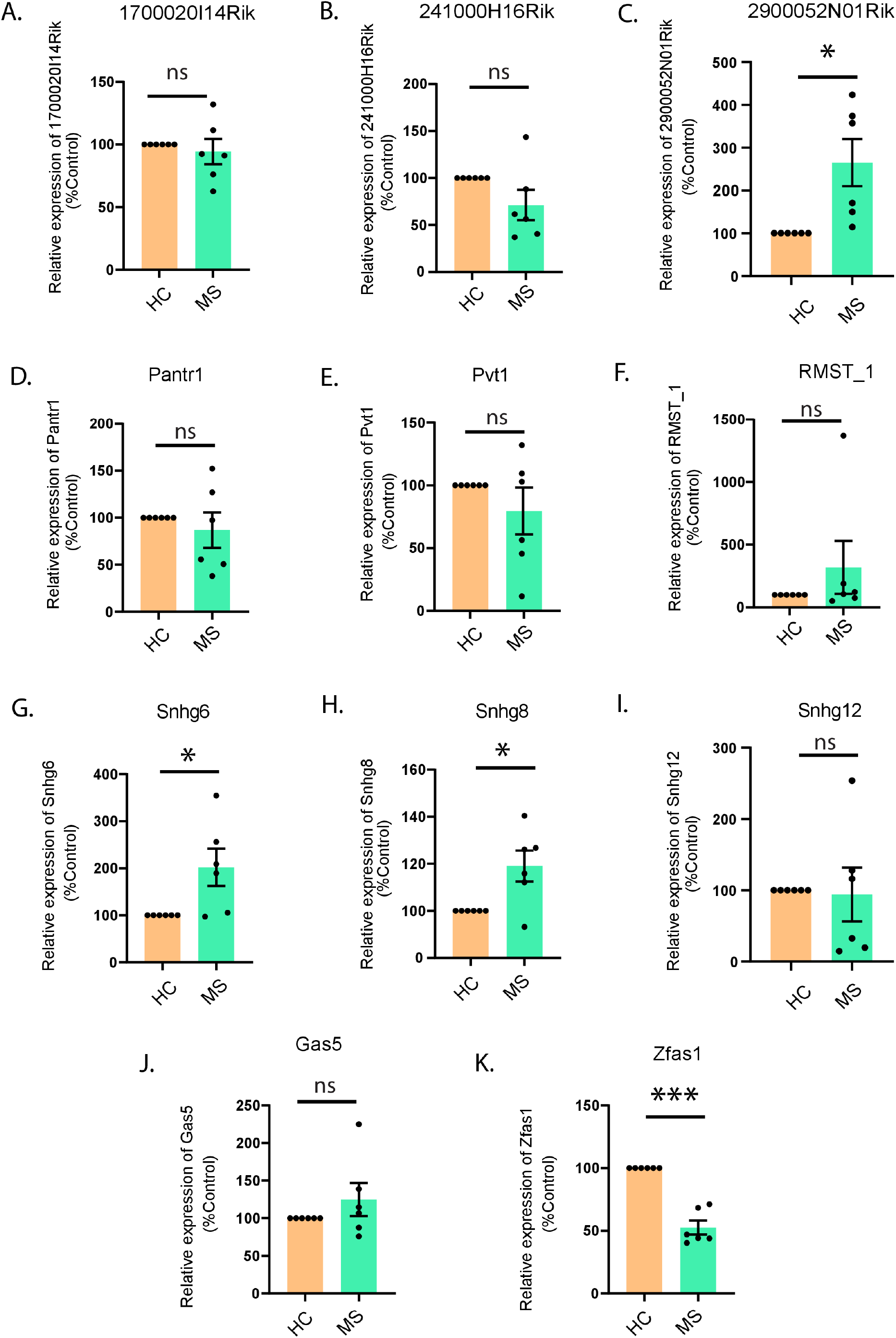
Maternal separation affects the expression of synapse-enriched lncRNAs in hippocampus of female P21 mice. Differential expression of synapse-enriched lnc-RNAs (A). 1700020I14Rik (B)2410006H16Rik, (C) 2900052NO1Rik, (D) Pantr1, (E) Pvt1, (F) RMST_1, (G) Snhg6, (H) Snhg8, (I) Snhg12, (J) Gas5, (K) Zfas1 as obtained through qPCR, *p < 0.05, **p < 0.01, ns = not significant. N=6, Data shown as Mean ± SEM. Unpaired Student’s t-test with Welch’s correction. The underlying data is also shared in https://figshare.com/articles/dataset/_b_Gas5_regulates_early_life_stress-induced_anxiety_and_spatial_memory_b_/26047969.

The *in situ* hybridization (Allen Brain Atlas) data showed that Gas5 is expressed in the CA1, CA2, CA3 and DG of both the dorsal and the ventral hippocampus (R. S. Patel et al. 2023), whereas, Pantr1 expression is mainly restricted to the CA1 region of the dorsal hippocampus (Goff et al. 2015). Differential expression of other lncRNAs is also observed in different neuronal populations of the hippocampus (CA1, CA2, CA3 and DG). For example, GM9968 is a lncRNA that is enriched in the hippocampus. Its expression spans through CA1, CA3 and DG in both the ventral and dorsal hippocampus, with a higher expression, especially in the ventral hippocampus. Meanwhile, another hippocampus-enriched lncRNA GM13644 is equally expressed in the ventral and dorsal hippocampus but higher at the DG compared to the CA1-CA3 regions (Kadakkuzha et al., 2015). These subregions of the hippocampus have been shown to have different functions (Tsien et al., 1996, Hitti and Siegelbaum, 2014, Nakashiba et al., 2008) and can also play a unique role in regulating the behaviour of an individual (Cui et al., 2022, Grinman et al., 2019, Butler et al., 2019). Existing literature shows that the dorsal hippocampus is involved in spatial memory formation (Moser and Moser 1998) and the ventral hippocampus has a prominent role in anxiety (Fanselow and Dong 2010). Therefore, to probe the lncRNA function in MS-induced anxiety and memory, we have focused on Gas5.

The Gas5 lncRNA is involved in aggression (Feldker et al. 2003) and novelty-induced behaviour (Xu et al. 2020). Consistent with these observations, our study demonstrates the involvement of Gas5 in MS-induced anxiety (Figure 3). The maternally separated mouse displays a significant reduction in exploring the displaced object in our spatial object recognition test, indicating deficits in memory upon early life stress (Figure 4). These memory deficits are partially rescued by the Gas5 knockdown (Figure 4). Our observation of impaired spatial learning following maternal separation is in agreement with previous findings (Huot et al. 2002)(Grace et al. 2009)(Hulshof et al. 2011). However, the impaired cognitive performance with accompanied enhancement in anxiety differs from other studies that demonstrated improved spatial learning along with elevated levels of anxiety following maternal separation (Suri et al. 2013) (McGrath, Records, and Rice 2008). This discrepancy could arise due to the use of different rodent strains and paradigms to test hippocampus-dependent memory. We speculate that heightened anxiety upon maternal separation could contribute to hyperactivity that may lead to the exploration of both the displaced as well as the non-displaced object in our memory test (Figure 4). Our data shows that there is no bias between home-caged and maternally separated mice in their preference towards any object. Also, no preferential bias for any object is observed between mice expressing Gas5 shRNA or non-targeting control (Figure 4).

**Figure 3.**
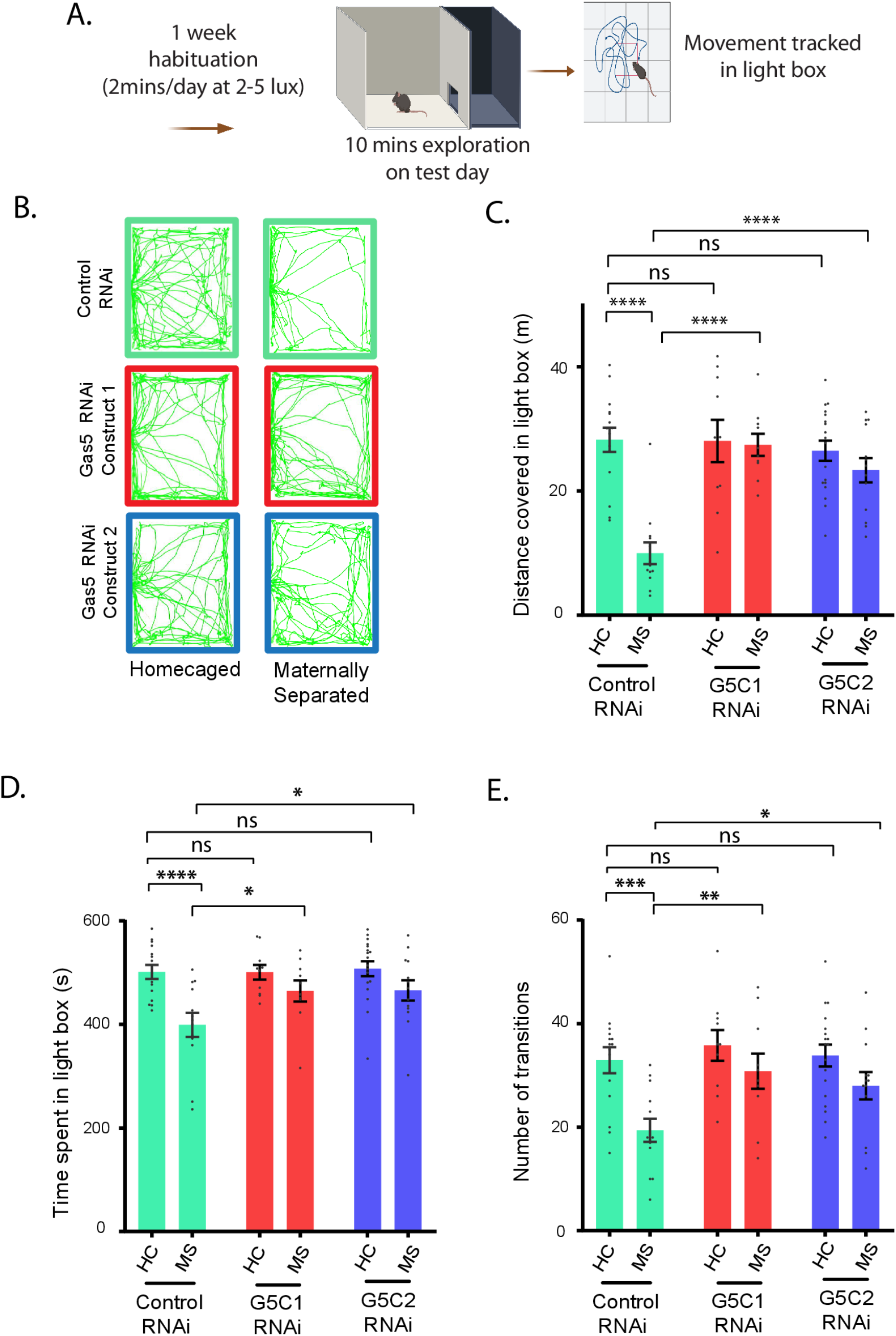
Gas5 knockdown reduces anxiety following maternal separation. **A**. Paradigm depicting light/dark box assay to measure anxiety. **B**. Representative trace showing the movement of the mouse in the bright-lit arena in the light/dark box. **C.** HC = Homecaged, MS = Maternally Separated, G5C1= shRNA # 1, G5C2 = shRNA # 2. Distance covered in the light box (in meters). **D.** Time spent in the light box (in seconds). **E.** Total number of transitions made between light and dark box. Data shown as mean±SEM, N=3-7, n=10 - 18, *p<0.05 **p< 0.01, ***p<0.005, ****p<0.0001, ns not significant. Two-way ANOVA with Fisher’s LSD test. The underlying data is also shared in https://figshare.com/articles/dataset/_b_Gas5_regulates_early_life_stress-induced_anxiety_and_spatial_memory_b_/26047969.

The role of direct Gas5 in hippocampus development is yet to be evaluated *in vivo,* although previous studies have shown that Gas5 has different expression patterns in young and old mice (Verbitsky et al., 2004). Aged mice showed a higher expression of Gas5. This finding was later supported by the fact that Alzheimer’s disease AD patients had an elevated level of Gas5 in their hippocampus accompanied by a decrease in hippocampal volume (Chen et al., 2022). The molecular mechanism of regulation of Gas5 in the development of the hippocampus is still unexplored. A study done in Mouse Embryonic Hippocampal Progenitor cells showed an elevated Gas5 level accompanies neuronal commitment, while Gas5 downregulation occurs during terminal differentiation, suggesting a regulatory role in conjunction with connexin expression during neurogenesis (Rozental et al., 2000).

It is important to note that the Gas5 RNAi alone does not influence anxiety or memory without any exposure to early life stress (Figure 3-4). This observation indicates that the MS-induced Gas5 level above the physiological set point is a prerequisite for anxiety and memory deficits due to early life stress. To evaluate whether the reversal of ES-induced anxiety and memory deficits occurs due to reduced level of Gas5 in the adult or developing brain, we have measured spine density and type from the adult hippocampus (P38 – P40) following shRNA-mediated knockdown of the transcript during development (P0). The spine density and type upon Gas5 knockdown were assessed as these parameters are key determinants of anxiety and memory (Melo et al., 2020; Leuner and Shors, 2013; Restivo et al., 2009; Kasai et al., 2010). No significant changes in the density of the total spine, stubby spine and mushroom spine were detected in the dendrites of hippocampal neurons (P38 - 40) following Gas5 knocked down (P0) (Figure 6). This observation confirms that the Gas5 activity during neuronal development does not influence the reversal of MS-induced anxiety and spatial memory deficit. The use of two distinct shRNAs against Gas5 (Figure S1) for all experiments eliminates the possibility of an off-target effect and consequent artefactual behavioural rescue or developmental changes. Taken together, our study highlights an RNA-centric approach that could potentially ameliorate the debilitating effect of early life stress.

**Figure 4.**
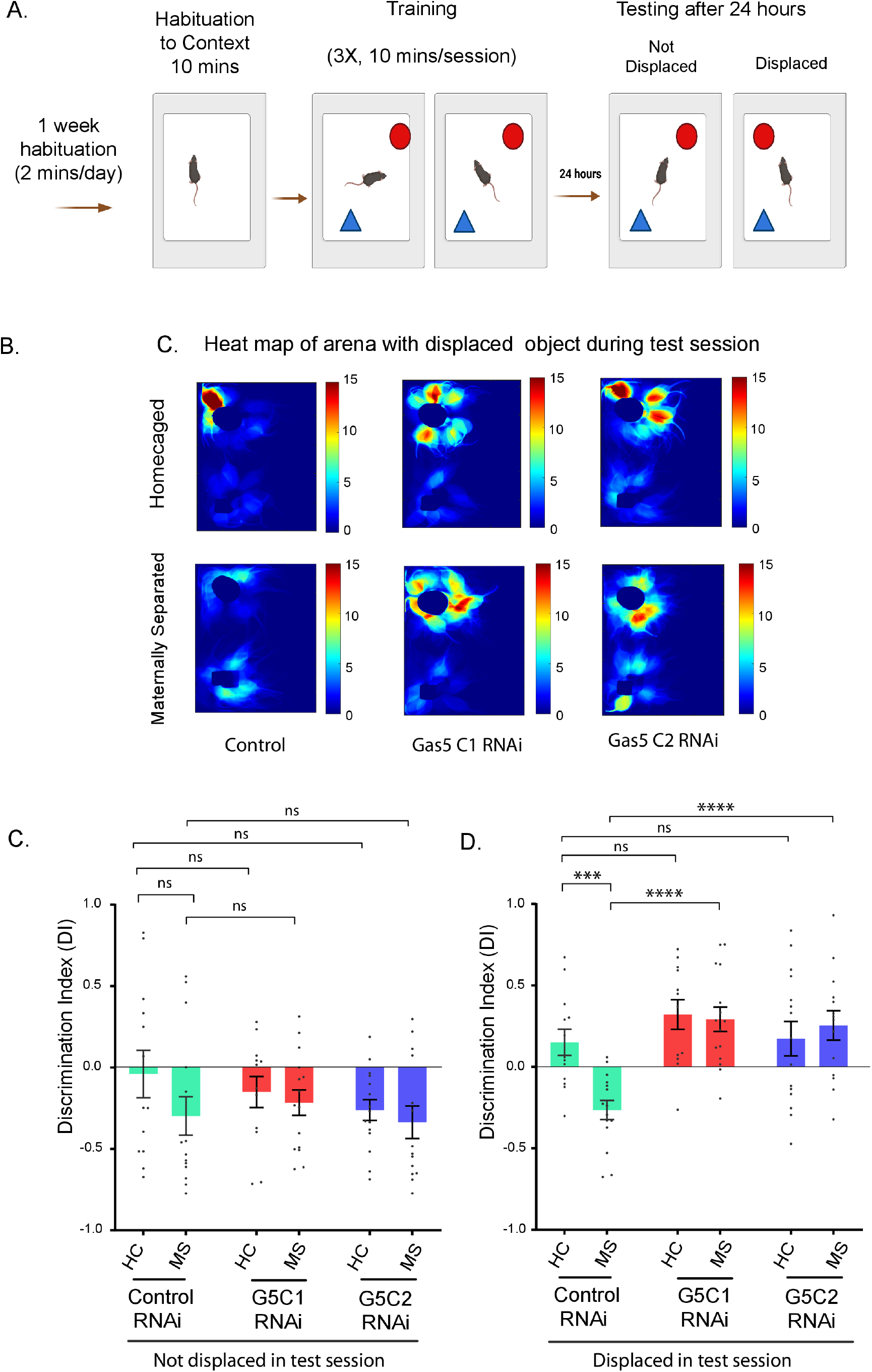
Gas5 reverses the effect of MS-induced deficits in memory. **A.** Schematic showing paradigm for SOR. **B.** Representative heat maps in the arena with the displaced object during the test session, circular (glass) object represents the displaced object. **C.** HC= Homecaged, MS= Maternally Separated, G5C1= shRNA # 1, G5C2= shRNA # 2. Discrimination Index (DI) in the arena with no displaced object during the test session. **D.** DI in the arena with a displaced object during the test session. Data shows mean±SEM. N=4-6, n=12-15, *p< 0.05, **p< 0.01, ***p<0.005, ns = not significant. Two-way ANOVA with Fisher’s LSD test. The underlying data is also shared in https://figshare.com/articles/dataset/_b_Gas5_regulates_early_life_stress-induced_anxiety_and_spatial_memory_b_/26047969.

**Figure 5.**
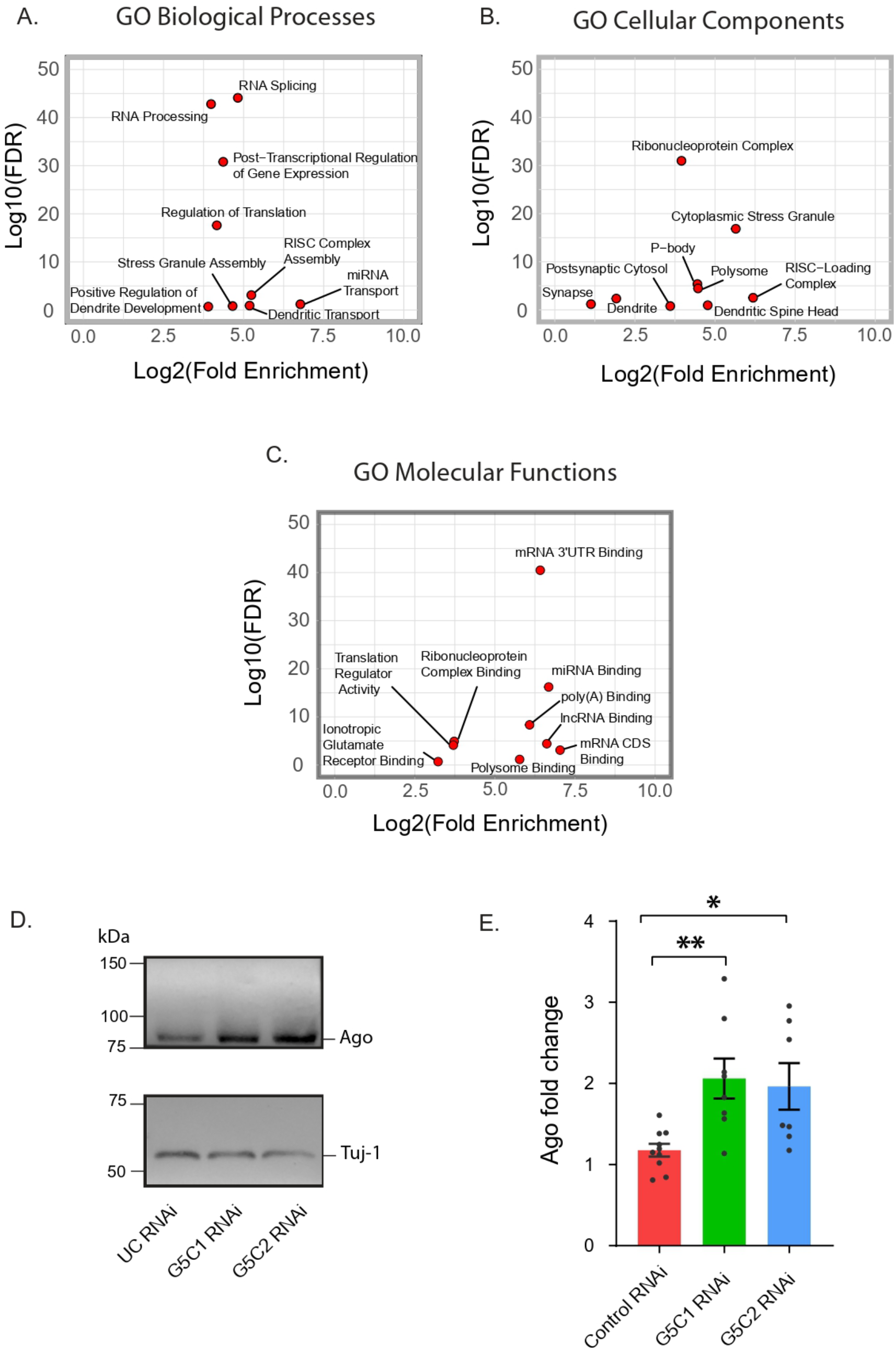
Involvement of Gas5 in RNA metabolism as revealed by GO-analysis. Available CLIP-seq data of Gas5 interacting coding transcripts were used to perform gene ontology (GO) analysis showing clusters of transcripts involved in **A**. Biological processes, **B**. Cellular components, and **C**. Molecular functions. Red dots show significant enrichment (FDR< 0.05), blue dots show highly enriched but not significant **D.** Immunoblot of Pan-Ago and Tuj-1 expression at the hippocampus of P21 male animals injected with Control (UC), Gas5 C1 (G5C1) or Gas5 C2 (G5C2) shRNA. **E.** Quantification of Pan-Ago with Tuj-1 as housekeeping gene; N=2-3, n=7-10, *p<0.05; Data shows mean±SEM. Unpaired t-test with Welch’s correction. The underlying data is also shared in https://figshare.com/articles/dataset/_b_Gas5_regulates_early_life_stress-induced_anxiety_and_spatial_memory_b_/26047969.

**Figure 6.**
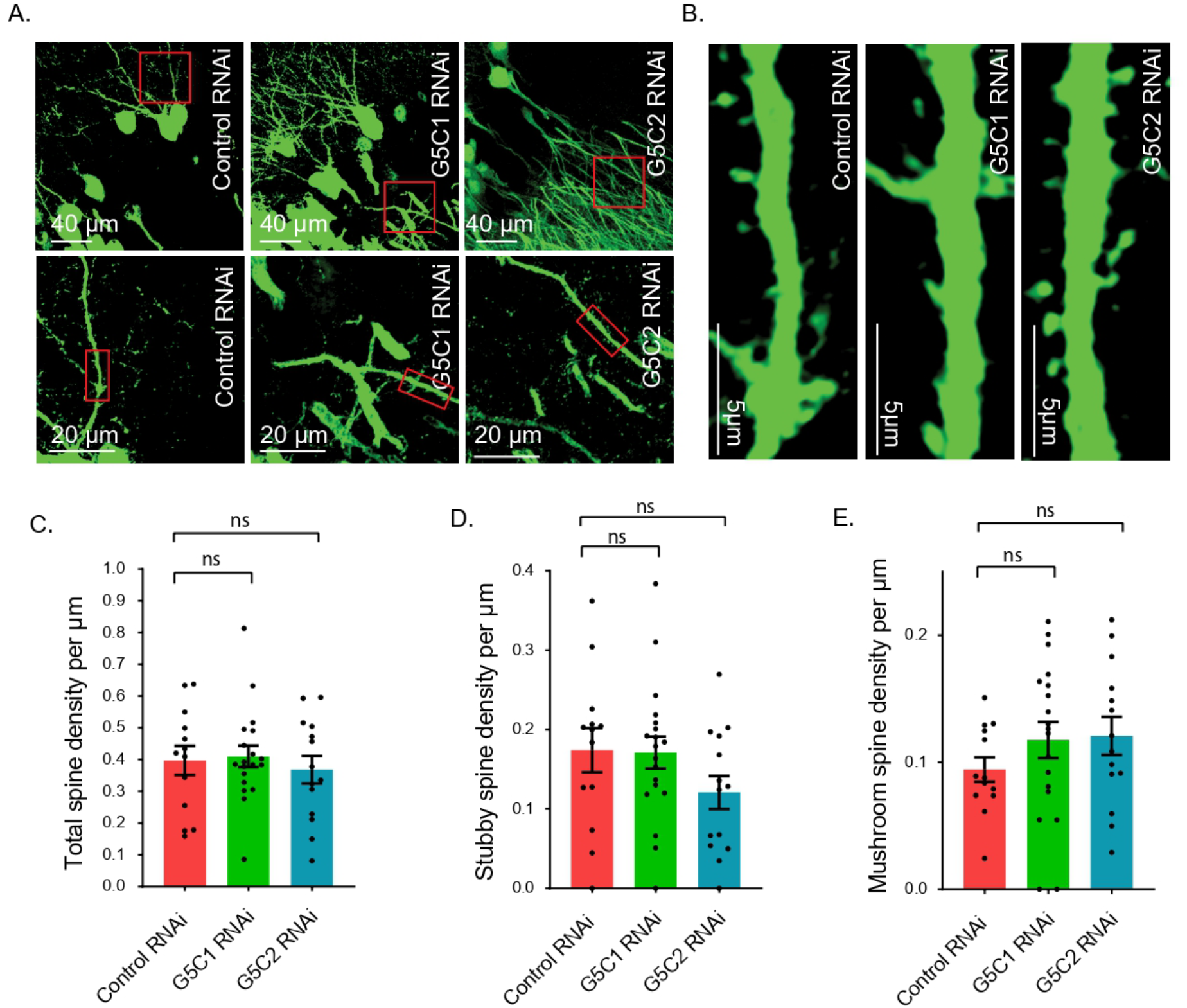
Effect of Gas5 knockdown on spine development. **A, B.** High magnification images of AAV injected mouse brain sections showing EGFP expressing hippocampal neurons; Control RNAi, G5C1 RNAi and G5C2 RNAi (left to right) respectively. **B**. Synapse density in hippocampal neurons of P30 mice injected with AAV at P0; Control RNAi, G5C1 RNAi and G5C2 RNAi (left to right) respectively. Quantification of **D.** total spine density, **E.** stubby spine density and **F.** mushroom spine density per µm in Control RNAi (UC), Gas5 C1 (G5C1) or Gas5 C2 (G5C2) shRNA; N=1, n=1-2, *n’*=13-19, *p<0.05; Data shows mean±SEM. Unpaired t-test with Welch’s correction. The underlying data is also shared in https://figshare.com/articles/dataset/_b_Gas5_regulates_early_life_stress-induced_anxiety_and_spatial_memory_b_/26047969.

For future investigations, employing alteration of Gas5 at distinct developmental stages (P7, P14, P21, P35, P50) would be an ideal choice to show the temporal resolution of Gas5 activity and its implications in reversing ES-induced elevated level of anxiety and memory deficit during adolescence. This region-specific knockdown of Gas5 through stereotaxic surgery, targeting specific regions such as CA1, CA2, CA3, and DG, could provide deeper insights into its function within the dorsal and ventral hippocampus. This can unravel the intricate role of Gas5 in modulating behaviour through tailored assays, shedding light on its potential therapeutic implications of Gas5. Further investigation is required to dissect a detailed mechanistic view of lncRNA-dependent reversal of elevated anxiety and memory deficit upon ES. To further understand the role of Gas5 in development, we can use the *in utero*-electroporation technique to knockdown Gas5 at the embryonic stage and thereafter study the development of the different types of the neural circuit of the hippocampus (i.e. Perforant pathway, Mossy fibre pathway, Schaffer collateral pathway etc.). To further understand the effect of Gas5 on functional changes at the synapse, we can study the changes in electrophysiological properties of neurons using the patch-clamp technique. This can help in elucidating the role of Gas5 in shaping the synaptic landscape of the hippocampus during crucial developmental periods.

To gain a plausible mechanistic insight into the Gas5-mediated regulation of anxiety and memory deficit upon maternal separation, we analyzed the functions of the Gas5 interacting transcripts identified from the POSTAR3 database. Our GO analysis revealed that Gas5 interacting transcripts are involved in RNA metabolism and synaptic functions. We have noted the regulation of transcripts associated with translation, RISC loading complex, miRNA transport and RNA splicing. The GO analysis also detected regulation of transcripts encoding proteins necessary for dendritic transport of ribonucleoprotein complex, somatodendritic compartment and ionotropic glutamate receptor binding. These observations impinge on the fact that Gas5 could, potentially, regulate synaptic function *via* post-transcriptional regulation of gene expression within subcellular spaces upon exposure to early life stress. Our data shows that the expression of Argonaute, a component necessary for miRISC loading and miRNA function, is reduced following the knockdown of Gas5. Consistent with previous observations showing the involvement of miRNAs in early life stress-induced gene expression (Allen and Dwivedi 2020), we speculate that Gas5-mediated regulation of anxiety and memory upon maternal separation involves components of the miRNA pathway. The enhanced expression of Argonaute may shift the miRNA loading between coding transcript and Gas5 *via* promoting molecular decoy activity to alter the translation status (Eulalio, Huntzinger, and Izaurralde 2008) (J. Li, Lv, and Che 2020). In our investigation, we identified six transcripts modulated by Gas5 (Fus, Hnrnph1, Hnrnpa2b1, Hnrnph2, Cpeb4 and Pcbp1), all pivotal in synaptic activity, using existing CLIP-seq data from the POSTAR3 database. Concurrently, other studies have focused on Gas5-mediated molecular framework regulating RNA processing operating in the infralimbic cortex for fear-memory extinction (Liau et al., 2023). The study proposes an interaction of Gas5 with Capirin1 (Cell Cycle Regulated Protein 1) and G3BP2 (Stress Granule Assembly Factor 2) at the synapse during fear extinction learning. The interaction of Gas5 with these RNA-granule proteins leads to the sequestering of Capirini and G3BP2 at the synapse. Caprin1 is evidenced to be important for the transport and translation of proteins essential for synaptic functions (Pavinato et al., 2023) and G3BP2 helps in RNA localization and stabilization (Jin et al., 2022). Therefore, sequestering of Caprin1 and G3BP2 alters synaptic plasticity through local translation of proteins. A separate study exploring Gas5’s role in addiction enlisted its targets in the nucleus accumbens (NcA), uncovering 16,053 potential interactors (Xu et al., 2020). Our comparative analysis revealed an overlap of 354 Gas5 targets between the NcA and hippocampus datasets, hinting at shared regulatory mechanisms across distinct brain regions.

We have performed a KEGG pathway and Gene Ontology analysis to determine the functional activities of these interacting partners of Gas5. The GO analysis revealed that Gas5 is predominantly involved in the regulation of RNA processing and translation of transcripts encoding proteins influencing synaptic function through the regulation of biological processes (pre and postsynaptic endocytosis, stress granule assembly, dendritic spine morphogenesis, regulation of axon extension, neuronal differentiation etc.), cellular components (polysome, growth cone, synapse, neuron projection etc.), and molecular functions (dopamine receptor binding, glutamate receptor binding, mRNA binding, GTPase activity etc.). KEGG pathway analysis indicated the involvement of Gas5 in key signalling pathways, c e.g. cAMP, mTOR and Wnt, that regulate synaptic functions in developing and adult brains (Ma et al., 2022, Grinman et al., 2021, Chen *et al*., 2020). These pathways have been shown to directly influence anxiety and memory (Gafford et al., 2013, Zhang et al., 2011, Moore and Loprinzi, 2021, Wang et al., 2020, Argyrousi et al., 2020, Bertagna et al., 2021) (Figure 7) For details see Table S2 and Supplementary data).

**Figure 7.**
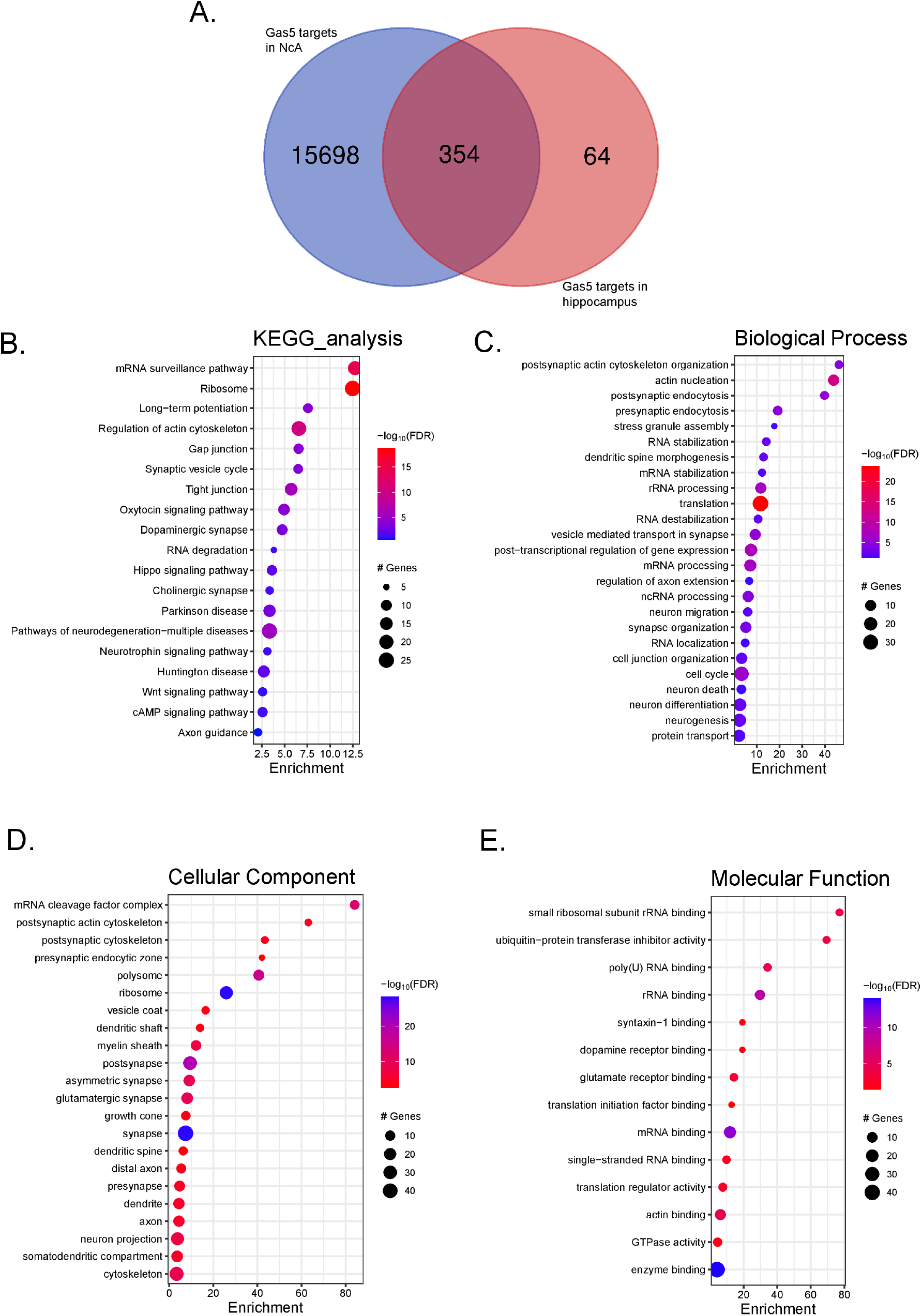
Gas5 targets are involved in the modulation of synapse and translation in the nucleus accumbens and the hippocampus. **A**. An overlap of the number of Gas5 targeted coding transcripts regulating addiction in the nucleus accumbens (NcA) (Xu et al. 2020) and fear memory extinction in the hippocampus (Liau et al. 2023). Gas5 interacting coding transcripts were used to analyse their involvement in the **B**. KEGG pathway and Enrichment pathways in Gene Ontology analysis of **C**. Biological processes, **D**. Cellular components, and **E.** Molecular functions. FDR<0.05. The underlying data is also shared in https://figshare.com/articles/dataset/_b_Gas5_regulates_early_life_stress-induced_anxiety_and_spatial_memory_b_/26047969.

In our study, the knockdown of synapse-enriched transcripts of Gas5 could lead to the sequestering of RNA-binding proteins (RBPs), disrupting translation dynamics at these neuronal junctions. The Gas5 transcript’s interaction with RBPs might influence how these proteins are compartmentalized within distinct synaptic subcellular spaces. This reorganization could restrict RBPs’ access to other mRNAs, causing shifts in local protein synthesis at the synapse and ultimately impacting synaptic plasticity.

Gas5 immunoprecipitation, coupled with Mass spectrometry analysis from hippocampal tissue of stressed animals, holds promise for uncovering novel protein candidates and pathways. By examining the varied expression of Gas5 targets, manipulating their levels via overexpression or loss of function can offer critical insights into the potential reversibility of early-life stress impacts. Leveraging CRISPR-Cas-inspired RNA targeting systems (CIRTs), Gas5 targets can be precisely targeted at the synapse, shedding light on their role in synapse development and plasticity. Additionally, it can also be used to rescue stress-induced anxiety and loss of memory offering a scope for therapeutic intervention.

The use of CO_2_ gas for euthanasia might be considered a limitation of our study as the high level of CO2 may trigger the expression of early-effect gene cFos and can also affect other gene expressions due to a panic attack at the moment of euthanasia (Johnson et al., 2011, Khokhlova et al., 2022). However, maternal separation data is normalised as the ratio between homecaged control and maternally separated animals. Hence, the probable artefact due to the euthanasia method is significantly minimized.

Overall, our study highlights the importance of evaluating diverse lncRNA activities during early life stress. To gain mechanistic insight, it would be important to understand how lncRNA activity is modulated by early life stress and its consequence on cognitive performance. We believe that this study will promote further research to probe how early life stress across temporal windows could lead to behavioural diversity *via* non-coding RNA-mediated regulation of gene expression.

## Supporting information

https://figshare.com/articles/dataset/_b_Gas5_regulates_early_life_stress-induced_anxiety_and_spatial_memory_b_/26047969

## AUTHOR CONTRIBUTION

D.B. and S.B. conceptualized the project. D.B. performed all experiments. S.S. performed spatial memory and anxiety analysis. D.B. and S.B. wrote the manuscript.

## ACKNOWLEDGEMENT

We thank Penn Vector Core for AAV vectors. We acknowledge Balakumar Srinivasan for GO analysis, Gourav Sharma for spatial memory set-up, Smruti Mhalgi for blind scoring of SOR data, Kamakshi Garg for assisting in confocal microscopy and Sarbani Samaddar for conceptual discussions and proofreading of this manuscript. This work is supported by extramural research grant BT/PR14071/GET/119/36/2015 (to S.B.) from the Department of Biotechnology, Government of India and the core fund of the National Brain Research Centre.

All art has been created with BioRender.com (Figure 1-4, S 1A-B, S 5 and Graphical Abstract).

## DATA AVAILABILITY

The data supporting this study’s findings are available at https://figshare.com/articles/dataset/_b_Gas5_regulates_early_life_stress-induced_anxiety_and_spatial_memory_b_/26047969. Additionally, a preprint of the manuscript is available on Biorxiv https://www.biorxiv.org/content/biorxiv/early/2023/09/27/2023.09.27.559778.full.pdf posted on 27th of September 2023.

## Abbreviations

Ago: Argonaute
AAV: Adeno-Associated Virus
AMPA: α-Amino-3-hydroxy-5-methyl-4-isoxazolepropionic acid
ANOVA: Analysis of Variance
BCA: Bicinchoninic Acid
BSA: Bovine Serum Albumin
CA1: Cornu Ammonis 1
CA2: Cornu Ammonis 2
CA3: Cornu Ammonis 3
CIRTs: CRISPR-Cas-inspired RNA targeting systems
CD1: Outbred Swiss mice
CLIP-seq: Cross-Linking Immunoprecipitation followed by high-throughput sequencing
CMV: Cytomegalovirus
CPCSEA: Committee for the Purpose of Control and Supervision of Experiments on Animals
CRISPR: Clustered Regularly Interspaced Short Palindromic Repeats
cDNA: Complementary DNA
DH5α: Escherichia coli strain
DI: Discrimination Index
DMEM: Dulbecco’s Modified Eagle Medium
DIV: Days In Vitro
DNA: Deoxyribonucleic Acid
dF: Degrees of Freedom
ECL: Enhanced Chemiluminescence
EDTA: Ethylenediaminetetraacetic Acid
EF1-α: Elongation Factor 1-alpha
EGFP: Enhanced Green Fluorescent Protein
ES: Early life stress
FBS: Fetal Bovine Serum
FDR: False Discovery Rate
Gas5: Growth Arrest Specific-5
GFP: Green Fluorescent Protein
GABAergic: Gamma-Aminobutyric Acidergic
GO: Gene Ontology
HEK293T: Human Embryonic Kidney 293 Cells, expressing large T antigen of SV40
HC: Homecaged
IAEC: Institutional Animal Ethics Committee
LSD: Least Significant Difference
MOI: Multiplicity of Infection
MRI: Magnetic Resonance Imaging
MS: Maternal separation
NcA: Nucleus accumbens
NEB: New England Biolabs
NMDA: N-Methyl-D-Aspartate
PBS: Phosphate Buffered Saline
PCR: Polymerase Chain Reaction
PFA: Paraformaldehyde
P: Postnatal Day
POSTAR3: Post-transcriptional Regulation Atlas of lncRNAs and mRNAs
Pvt1: Plasmacytoma Variant Translocation 1
qPCR: Quantitative Polymerase Chain Reaction
RBPs: Ribosome Binding Proteins
qRT-PCR: Quantitative Reverse Transcription Polymerase Chain Reaction
RISC: RNA-Induced Silencing Complex
Rmst1: rhabdomyosarcoma 2-associated transcript 1
RNAs: Ribonucleic acids
RNP: Ribonucleoprotein
SDS: Sodium Dodecyl Sulfate
SEM: Standard Error of the Mean
shRNAs: Short hairpin RNAs
Snhg8: Small Nucleolar RNA Host Gene 8
Snhg12: Small Nucleolar RNA Host Gene 12
SV40: Simian Virus 40
SynLAMP: Synaptic Lysosome-associated Membrane Protein
Tubulinβ-1: Beta-1 tubulin
UC: Universal Control
Zfas1: Zinc finger antisense 1
ZsGreen: Zoanthus sp. Green Fluorescent Protein

