## Supplementary material for "Gas5 regulates early life stress-induced anxiety and spatial memory": https://figshare.com/articles/dataset/_b_Gas5_regulates_early_life_stress-induced_anxiety_and_spatial_memory_b_/26047969

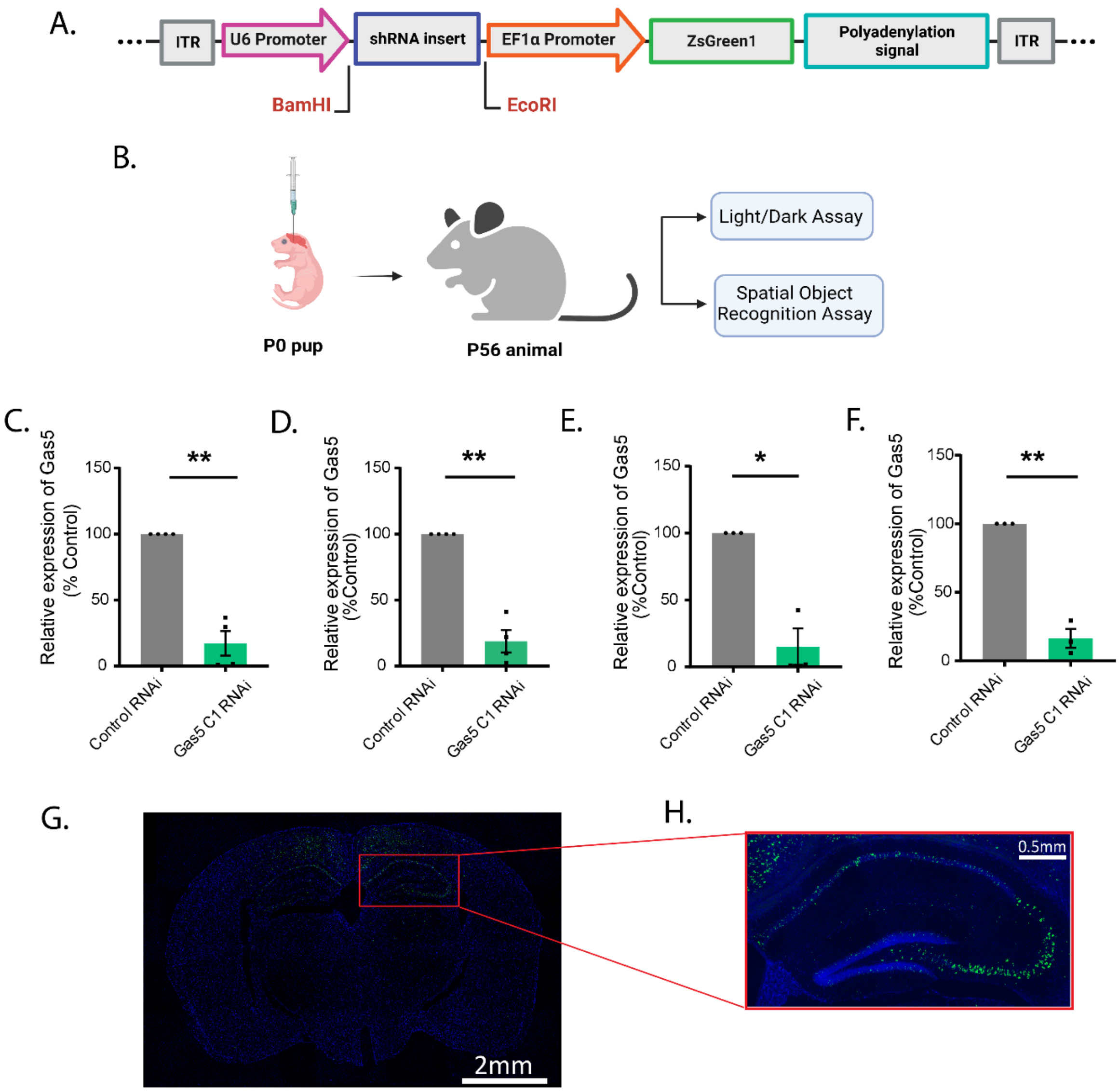

A.
ITR
U6 Promoter
shRNA insert
EF1α Promoter
ZsGreen1
Polyadenylation signal
ITR
BamHI
EcoRI
B.
P0 pup
P56 animal
Light/Dark Assay
Spatial Object Recognition Assay
C.
Relative expression of Gas5 (% Control)
150
100
50
0
**
Control RNAi
Gas5 C1 RNAi
D.
Relative expression of Gas5 (%Control)
150
100
50
0
**
Control RNAi
Gas5 C1 RNAi
E.
Relative expression of Gas5 (%Control)
150
100
50
0
*
Control RNAi
Gas5 C1 RNAi
F.
Relative expression of Gas5 (%Control)
150
100
50
0
**
Control RNAi
Gas5 C1 RNAi
G.
2mm
H.
0.5mm

**Figure S1. Gas5 knockdown by stereotaxic assay in P0 pups**. **A**. Map of vector used for Gas5 knockdown where shRNA is expressed under U6 promoter and Zs-Green under EF1-α; **B**. Stereotaxic injection of AAV was done in P0 pups which were later used for behavioural assays, like light/dark box assay and spatial object recognition assay, at P56. **C**. Gas5 RNAi tested in primary neuronal culture: G5C1 RNAi with GAPDH as internal control, N=4, **D.** with Tubulin-β1 as internal control, N=4, **E.** G5C2 RNAi with GAPDH as internal control, N=3, **F.** with Tubulin-β1 as internal control, N=3. $*p < 0.05$, $**p < 0.01$. Data shown as mean±SEM. Paired student's t-test. **G-H**. Brain section of P58 mouse showing expression of AAV inducing Gas5 knockdown in hippocampi. **G.** whole brain section (Leica fluorescence microscope LAS V4.2) and **H**. hippocampus magnified (Zeiss apotome). *(Please see Figures 3 and 4 for behaviour data after viral infection).*

The underlying data is also shared in https://figshare.com/articles/dataset/_b_Gas5_regulates_early_life_stress-induced_anxiety_and_spatial_memory_b_/26047969 .

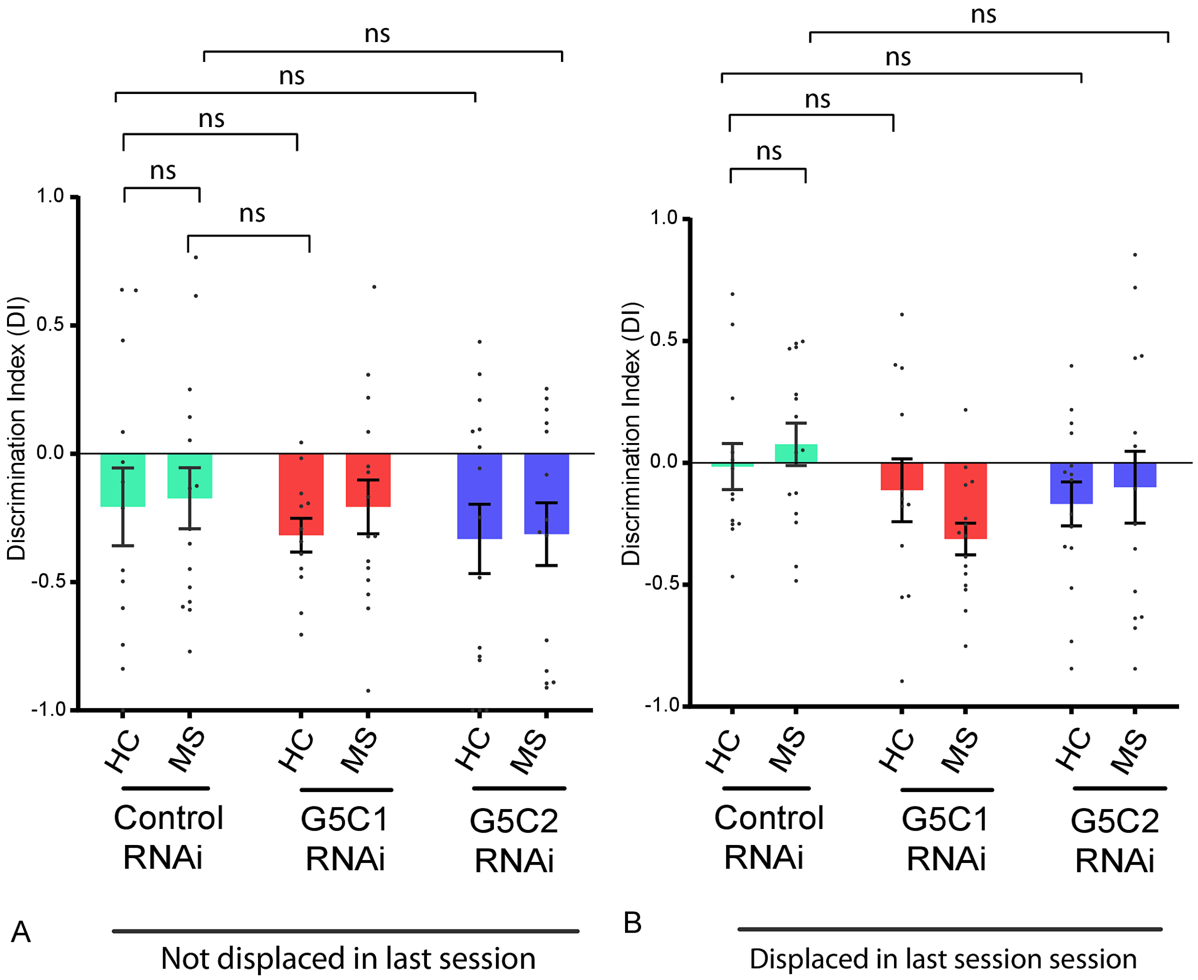

ns
ns
ns
ns
ns
1.0
0.5
0.0
-0.5
-1.0
Discrimination Index (DI)
HC
MS
HC
MS
HC
MS
Control RNAi
G5C1 RNAi
G5C2 RNAi
A
Not displaced in last session
ns
ns
ns
ns
1.0
0.5
0.0
-0.5
-1.0
Discrimination Index (DI)
HC
MS
HC
MS
HC
MS
Control RNAi
G5C1 RNAi
G5C2 RNAi
B
Displaced in last session session

**Figure S2. Discrimination index of animals during training. A**. Discrimination Index from the last session in the arena where no object was displaced during the test session, **B**. Discrimination Index from the last session in the arena where an object was displaced during the test session. Data shows mean±SEM. Two-way ANOVA with Fisher's LSD, N=4-6, n=12-15, ns = not significant. *(Please see Figure 4 also).*

The underlying data is also shared in https://figshare.com/articles/dataset/_b_Gas5_regulates_early_life_stress-induced_anxiety_and_spatial_memory_b_/26047969 .

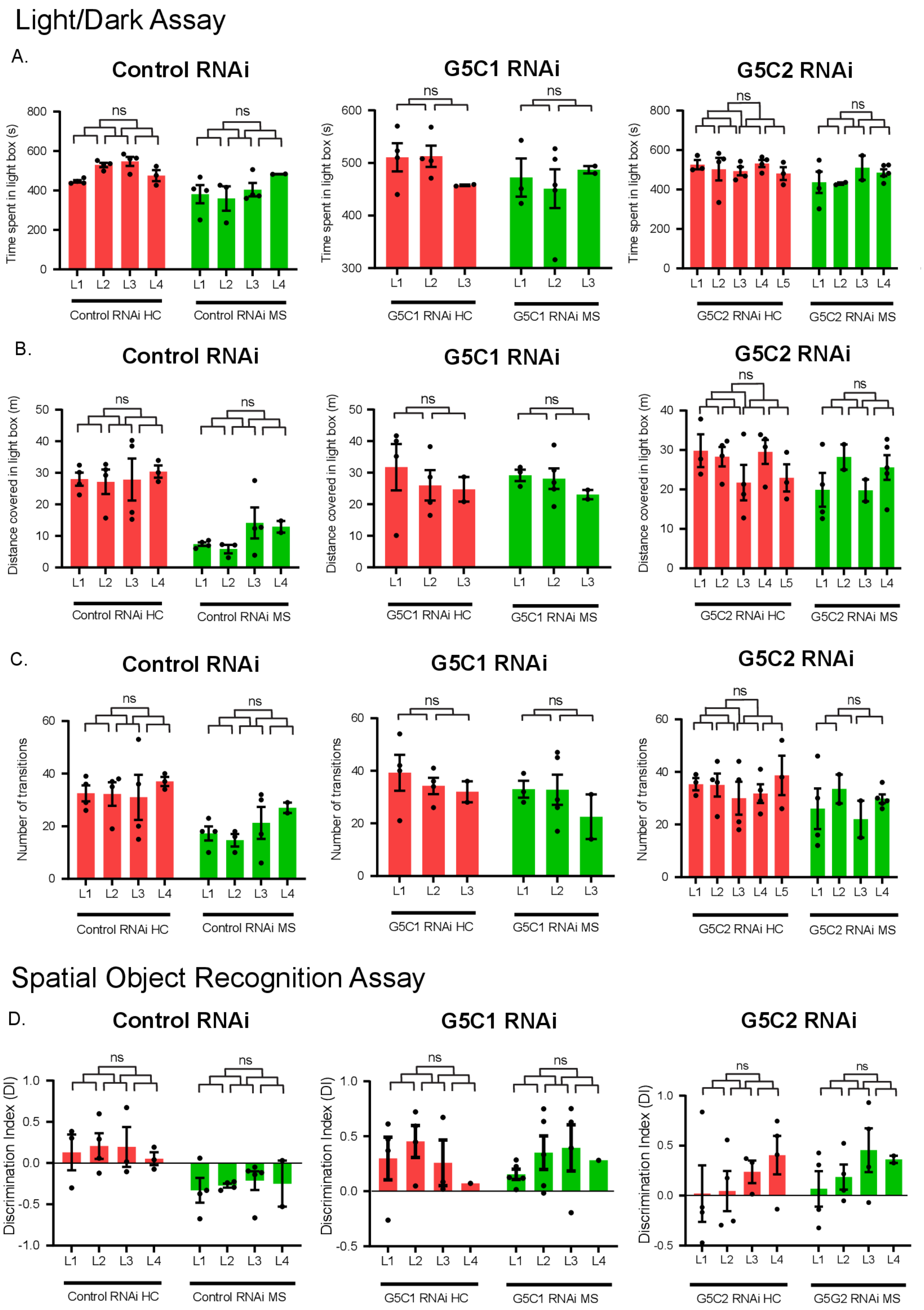

Light/Dark Assay
A.
Control RNAi
G5C1 RNAi
G5C2 RNAi
Time spent in light box (s)
ns
L1 L2 L3 L4
L1 L2 L3 L4 L5
Control RNAi HC
Control RNAi MS
G5C1 RNAi HC
G5C1 RNAi MS
G5C2 RNAi HC
G5C2 RNAi MS
B.
Control RNAi
G5C1 RNAi
G5C2 RNAi
Distance covered in light box (m)
C.
Control RNAi
G5C1 RNAi
G5C2 RNAi
Number of transitions
Spatial Object Recognition Assay
D.
Control RNAi
G5C1 RNAi
G5C2 RNAi
Discrimination Index (DI)
G5G2 RNAi MS

**Figure S3. Litter effect amongst animals in all behavioural experiments. A-C**. Litter effect of animals used for light dark assay with respect to **A.** Time, **B.** Distance, **C**. Transitions in Control RNAi, G5C1 RNAi and G5C2 RNAi animals of HC and MS conditions. Data shown as Data shows mean±SEM. Two-way ANOVA with Fisher's LSD, N= 3-5, n=2-5, ns = not significant. **D**. Litter effect of animals used for spatial object recognition assay, Control RNAi, G5C1 RNAi and G5C2 RNAi animals of HC and MS conditions. Data shown as Data shows mean±SEM. Two-way ANOVA with Turkey's correction, N= 4, n=1-5, ns = not significant. *(Please see Figures 3 and 4 also).*

The underlying data is also shared in https://figshare.com/articles/dataset/_b_Gas5_regulates_early_life_stress-induced_anxiety_and_spatial_memory_b_/26047969 .

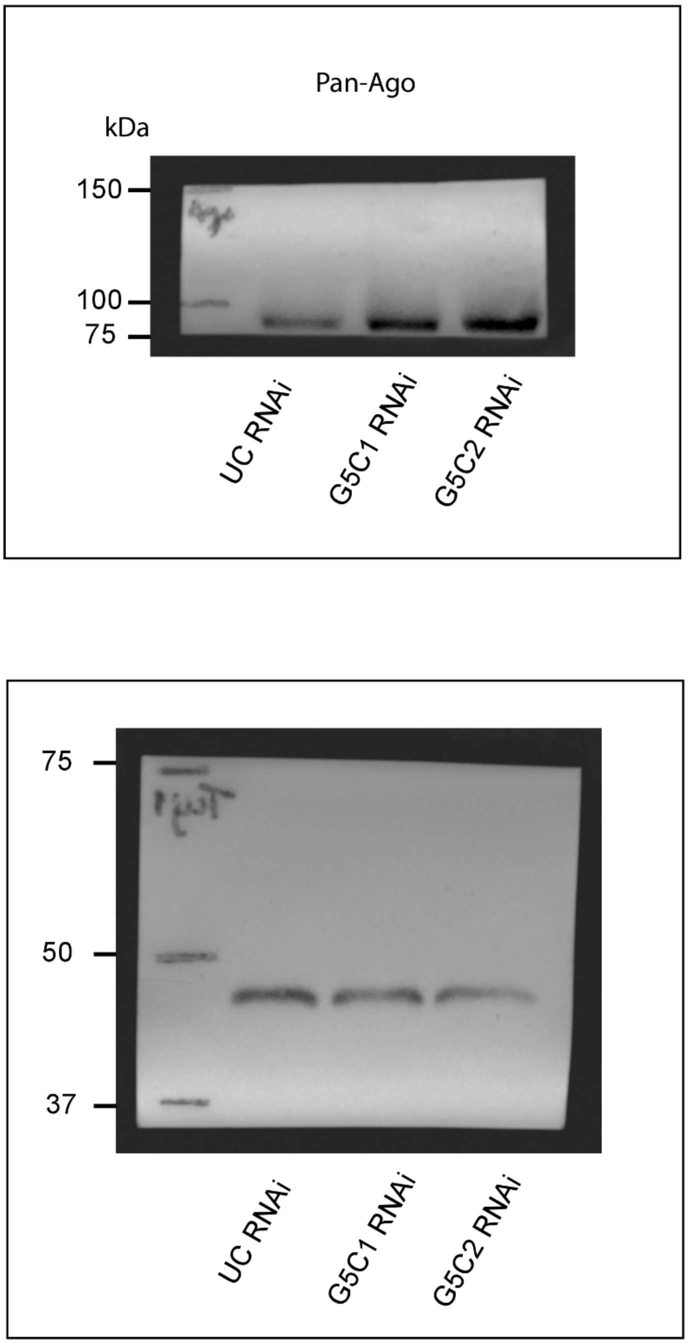


**Figure S4. Full immunoblot** of **A.** Pan-Ago and **B.** Tuj-1 expression at the hippocampus of P21 male animals injected with Control (UC), Gas5 C1 (G5C1) or Gas5 C2 (G5C2) shRNA. *(Please see Figure 5D also).*

The underlying data is also shared in [https://figshare.com/articles/dataset/_b_Gas5_regulates_early_life_stress-induced_anxiety_and_spatial_memory_b_/26047969](https://figshare.com/articles/dataset/_b_Gas5_regulates_early_life_stress-induced_anxiety_and_spatial_memory_b_/26047969) .

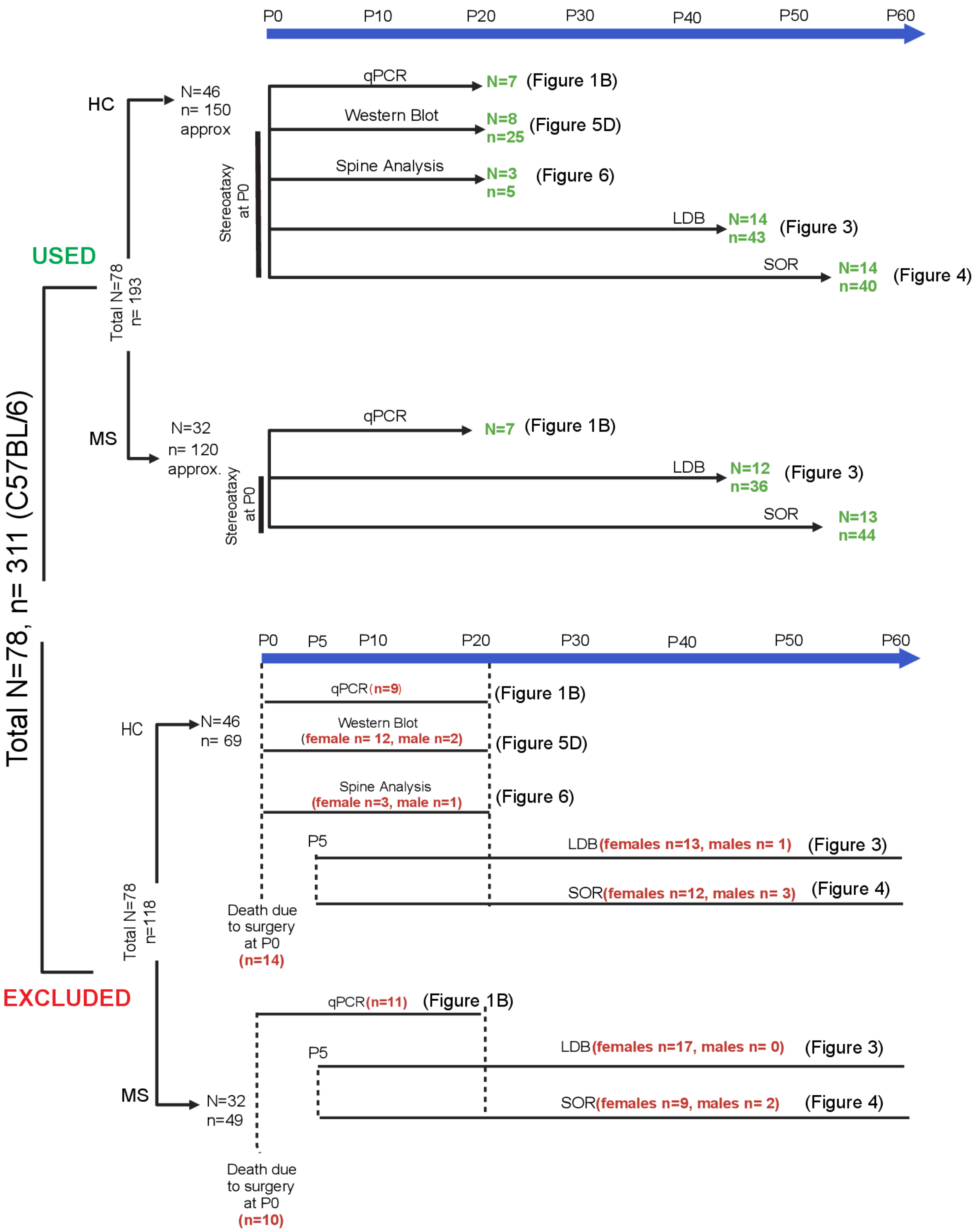


Total N=4, n= 30 approx. (CD1) → Dam was sacrifice and pup brains (hippocampus) were pooled for culture

**Figure S5. Time diagram of the number of animals used for different experiments and time points of exclusion of animals**. N= number of pregnant dams/number of litters, n=number of pups. Green = animals used in experiments, Red = animals excluded from the experiment at various points. All stereotaxic surgeries were performed at P0 (total 24 pup deaths due to surgery were recorded between P0-P5 across all experiments: Spine analysis, Light/dark box assay, Spatial object recognition assay). P21 pups from pregnant dams (N=14 (HC+MS)) were used for qPCR **(Figure 1B, 2).** 20 pup deaths were recorded between P0-P21 in animals used for qPCR. Brains were pooled from the surviving pups for qPCR (in both males and females in HC and MS conditions). N=26, n=79 animals were used for light/dark assay **(Figure 3**), death of 1 male was recorded (P0-P60), and total 30 female pups died between P0-P21 or were sacrificed at P21. N=27, n=84 animals were used for spatial object recognition assay **(Figure 4)**, death of 5 male pups was recorded (P0-P60), and total 21 female pups died between P0-P21 or were sacrificed at P21. N=8, n=25 male animals were used for the Western blot experiment **(Figure 5)**, the death of 2 male pups was recorded (between P6-P21) and total 12 female pups died between P0-P21 or were sacrificed at P21. N=3, n=5 male animals were used for the Spine analysis **(Figure 6)**, the death of 1 male pup was recorded (P0-P21) and total 3 female pups died between P0-P21 or were sacrificed at P21.
